# A weak coupling mechanism mediates the recovery stroke of myosin VI: a free energy simulation and string method analysis

**DOI:** 10.1101/2023.10.02.560423

**Authors:** Florian E.C. Blanc, Anne Houdusse, Marco Cecchini

## Abstract

Myosin motors use the energy of ATP to produce force and directed movement on actin by a swing of the lever arm. ATP is hydrolysed during the off-actin re-priming transition termed recovery stroke. To provide an understanding of chemo-mechanical transduction by myosin, it is critical to determine how the reverse swing of the lever arm and ATP hydrolysis are coupled. Previous studies concluded that the recovery stroke of myosin II is initiated by closure of the Switch II loop in the nucleotide-binding site. Recently, we proposed that the recovery stroke of myosin VI starts with the spontaneous re-priming of the converter domain to a putative pre-transition state (PTS) intermediate that precedes Switch II closing and ATPase activation. Here, we investigate the transition from the pre-recovery, post-rigor (PR) state to PTS in myosin VI using geometric free energy simulations and the string method. First, our calculations rediscover the PTS state agnostically and show that it is accessible from PR via a low free energy transition path. Second, separate path calculations using the string method illuminate the mechanism of the PR to PTS transition with atomic resolution. In this mechanism, the initiating event is a large movement of the converter/lever-arm region that triggers rearrangements in the Relay-SH1 region and the formation of the kink in the Relay helix with no coupling to the active site. Analysis of the free-energy barriers along the path suggests that the converter-initiated mechanism is much faster that the one initiated by Switch II closure, which supports the biological relevance of PTS as a major on-pathway intermediate of the recovery stroke in myosin VI. Our analysis suggests that lever-arm re-priming and ATP hydrolysis are only weakly coupled, so that the myosin recovery stroke is mostly driven by thermal fluctuations and stabilised by nucleotide consumption via a ratchet-like mechanism.

**Author summary:** Myosin is an ATP-powered motor protein that is crucial for cellular functions like muscle contraction, cell division, and the transport of molecular cargos. Understanding how myosin transforms chemical energy from ATP hydrolysis into mechanical work is a central open question in structural bioenergetics, which could guide the design of next-generation synthetic nanomachines. In myosin, ATP hydrolysis is coupled to the re-priming of the lever-arm during the recovery stroke transition. Using advanced molecular dynamics simulations, we described the sequence of events and the energetics of the recovery stroke of myosin VI with an unprecedented level of detail. The results support a mechanism in which the re-priming of the mechanical amplifier region of the protein is almost entirely driven by thermal fluctuations and stabilized by ATP hydrolysis. This weakly-coupled mechanism suggests that myosin can “rectify” thermal fluctuations to work efficiently in an isothermal environment dominated by stochastic fluctuations like the cell.

## Introduction

Myosins are a superfamily of actin-based motor proteins involved in cellular functions such as cargo transport, endocytosis, cell division and motility [1]. Force and directional movement are produced upon strong binding of the motor domain to F-actin, which triggers the sequential release of the ATP hydrolysis products and the forward movement of the lever-arm domain, or powerstroke [1–3], see Fig 1a. Binding of a new ATP molecule promotes detachment from F-actin and initiates the recovery stroke, *i.e.*, the transition during which the lever-arm is re-primed to its armed configuration. In absence of actin, this transition is rather fast (≈1 ms) and reversible. Interestingly, ATP is hydrolyzed at the end of the recovery stroke rather than during the powerstroke. Therefore, the mechanistic elucidation of this complex, ∼1 ms conformational transition is necessary to understand the structural basis for chemo-mechanical transduction in myosin. Myosin VI (Myo6) is the only-known minus-directed motor, and fulfills specific cellular roles [4, 5]. Myo6 harbors two unique inserts including one responsible for directionality reversal, but otherwise shares the same design plan as other myosins [6–8].

**Fig 1.**
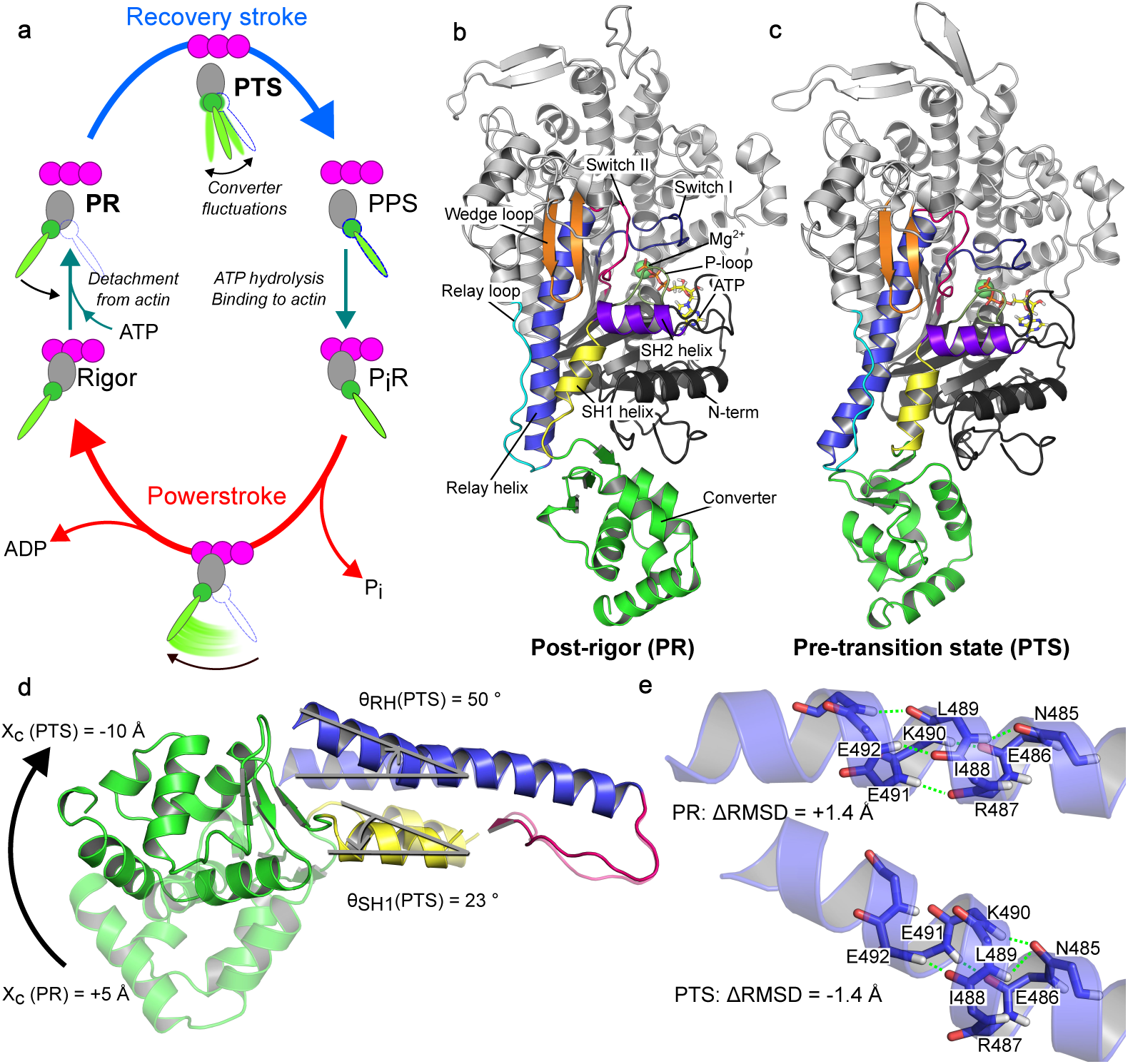
Overview of the PR → PTS transition. (a) Simplified Myosin VI functional cycle including the putative PTS. (b) Equilibrated PR structure highlighting the structural elements involved in the recovery stroke. (c) Equilibrated PTS structure, colored as in b. (d) Comparison between PR (transparent) and PTS (opaque) showing the movement of the converter, the tilting of the SH1 helix, and the kink and re-orientation of the C-terminal fragment of the Relay Helix. Typical values for the corresponding CVs are given. (e) Comparison of the backbone structure of residues 485-491 of the Relay helix before and after kink formation. Typical values of the ΔRMSD are given.

High-resolution crystal structures of the initial state (post-rigor state, PR) and final state (pre-powerstroke state, PPS) of the recovery stroke of *Dictyostelium discoideum* Myosin 2 (Dd Myo2) as well as Myo6 have revealed the main subdomain rearrangements, which are similar for both motors. Chief among them are: the re-priming of the converter/lever-arm; the formation of a kink halfway along the conserved Relay Helix (RH); and the closure of the Switch II loop over the *γ*-phosphate of ATP [7, 9]. This latter rearrangement turns on ATPase function by stabilizing an active site geometry conducive to hydrolysis [10–13]. A key open question is how Switch II closure couples to converter re-priming. To date, most computational analyses of the PR→PPS transition mechanism have predicted that Switch II closure initiates the recovery stroke [14–25]. Among them, the popular model by Fischer and co-workers posits a strongly, or *mechanically* coupled mechanism where Switch II closure directly drives the rotation of the converter [14, 17]. In this model, structural changes occur in progressive, concerted fashion and involve a two-step rearrangement in the RH, which sits between the active site and the converter.

In sharp contrast, we have recently proposed a mechanism for the recovery stroke of Myo6 in which the rotation of the converter precedes the closure of Switch II and is uncoupled from it until the later stages of the transition [26]. Because this mechanism entails near complete converter re-priming driven purely by random fluctuations, we named it the ”ratchet-like” model. It involves a putative structural intermediate called pretransition state (PTS), which we characterized by X-ray crystallography and all-atom Molecular Dynamics (MD), see Fig 1a. The PTS structure (PDB: 5O2L) exhibits hallmarks of the post-recovery state, namely a (nearly) re-primed converter and a kink in the Relay helix (Figure 1b-e), yet with an open Switch II. The ratchet-like model of the recovery stroke therefore comprises two major conformational transitions, the PR → PTS transition in which (most of) the converter re-priming takes place, and the PTS → PPS transition in which Switch II closes. In this scenario, converter swing is weakly, or *statistically*, coupled to Switch II closure.

Whether the recovery stroke of Myo6 proceeds by a strongly-coupled, Switch II-initiated or statistically-coupled, ratchet-like mechanism remains an open question. To explore the recovery stroke mechanism with atomic resolution, we turn to all-atom MD simulations with enhanced sampling strategies. Geometric free energy calculations (including the adaptive biasing force method, or ABF) evaluate the potential of mean force (PMF) along low-dimensional collective variables (CVs) describing the transition [27–30]. The related string method in collective variables (CVSM) identifies the minimum free energy path (MFEP) in a CV-space of arbitrary dimension [31, 32], which makes it ideal to elucidate the transition mechanism between two known structures. The string method has been used to describe conformational transitions in molecular machines, see for example [33–35]. For myosin, Ovchinnikov et al. applied it to the internal conformational transition of the Myo6 converter [36, 37] and Cui et al. used it in combination with a quantum potential to map ATP hydrolysis pathways in Dd Myo2 [12]. To our knowledge, the string method has not been applied yet to the elucidation of conformational transitions of the full-length myosin motor domain.

We report here on a computational analysis of the mechanism and energetics of the transition from PR to PTS, *i.e.*, the first major step of the recovery stroke in the ratchet-like model. First, we use ABF to map a coarse-grained free energy landscape of the transition. Consistent with the ratchet-like prediction, we identify agnostically a state corresponding to PTS, and show that it is accessible from PR by a low free energy path, supporting its relevance as an on-pathway intermediate. Second, we perform path optimizations with the CVSM to determine a higher-resolution description of the transition. These independent calculations agree remarkably well with the ABF map and reveal a plausible mechanism by which near-complete re-priming of the mechanical amplifier region is triggered by a spontaneous converter/lever-arm swing, eventually resulting in the formation of the kink in the Relay helix while Switch II remains open. Despite some uncertainty as to the nature of the rate-limiting step (lever-arm movement vs formation of the kink), free energy barriers suggest that the ratchet-like pathway is explored much faster than the Switch-II initiated one. Therefore, our results support the PTS state as a functional intermediate of the myosin VI cycle, and the ratchet-like model for the myosin VI recovery stroke.

## Results

To investigate the recovery stroke of Myo6, we combine extended ABF (eABF, [38]) and the CVSM to take advantage of their respective strengths. We introduce CVs *X_c_* (Fig. 1d) to measure the position of the converter relative to the motor domain and Δ*RMSD* (Fig. 1e) to describe the local conformation of the RH (*i.e.,* the formation of the kink); see Methods for a precise definition. Together, these CVs describe two characteristic rearrangements from PR to PTS. We perform an eABF calculation over (*X_c_,* Δ*RMSD*) to determine the coarse-grained free energy landscape of early recovery stroke in the ratchet-like model, starting from the PR state. Then, we carry out string optimizations in a higher-dimensional space encompassing degrees of freedom left out of the eABF calculation, to determine a high-resolution picture of the transition mechanism from PR to PTS. This strategy yields an integrated structural and energetic description of the early stages of the recovery stroke and enables consistency checks between separate sets of simulations.

### Coarse-grained free energy landscape of the PR → PTS transition

The full coverage of configurational space, smooth convergence of the free energy gradient estimates and low estimated statistical error suggest that the eABF calculations are converged (see Supplementary Text 1).

### eABF discovers the PTS state without prior knowledge

The free energy landscape computed by eABF reveals 5 local free energy minima, including the PR basin (upper right corner), see Fig. 2a. Strikingly, a wide basin encompassing 2 sub-minima is found in the lower left corner of the free energy landscape, corresponding to a kinked Relay helix and (partially) re-primed converter. Interestingly, we observe that the active site -as characterized by the *d*_1_ and *d_γ_* distances, see Methods-remains open throughout eABF sampling, including in the identified metastable states (Supplementary Figure S4a). Therefore, the calculations indicate that there exists a metastable Myo6 configuration with open active site, nearly re-primed converter and kinked Relay helix. We note that if the closure of Switch II were strongly coupled to converter movement and/or Relay helix kink formation, closed active-site conformers would have been preferentially sampled in the lower-left region of the free energy landscape, which is not the case. Additionally, this basin’s characteristics are consistent with the PTS state, as evidenced by comparison to previously reported unbiased MD simulations of Myo6 [26], see Supplementary Fig. S4b. We find that both sub-basins (labelled PTS1 and PTS2 in Fig 2a) correspond to converter positions and Relay helix conformations explored in unbiased PTS simulations. Finally, representative conformers extracted from the PTS1 basin by clustering demonstrate marked structural similarity to the PTS crystal structure (Fig. 2b). Taken together, these findings imply that not only a partially re-primed state with open active site exists, but also that the PTS structure is representative of this state, indicating that eABF has discovered the PTS state without prior knowledge of the PTS structure (see Methods). The finer structure of the free energy landscape suggests -also in agreement with earlier unbiased MD [26]-that the PTS state encompasses at least two distinct metastable states, PTS1 and PTS2, separated by a low, 3 kcal mol^−1^ free energy barrier, and which correspond to slightly different RH conformers and average converter positions. The PTS2 converter position is closer to PPS, suggesting it is on the recovery stroke pathway. From PTS1, the transition to PTS2 entails a further 7 Å converter movement associated with a slight re-orientation of the RH. This transition corresponds to rapid fluctuations within the PTS basin, as demonstrated by their reversible sampling on the ∼100 ns timescale in earlier unbiased MD simulations, and agrees with our previous observation that the converter is highly dynamic in the PTS state [26].

**Fig 2.**
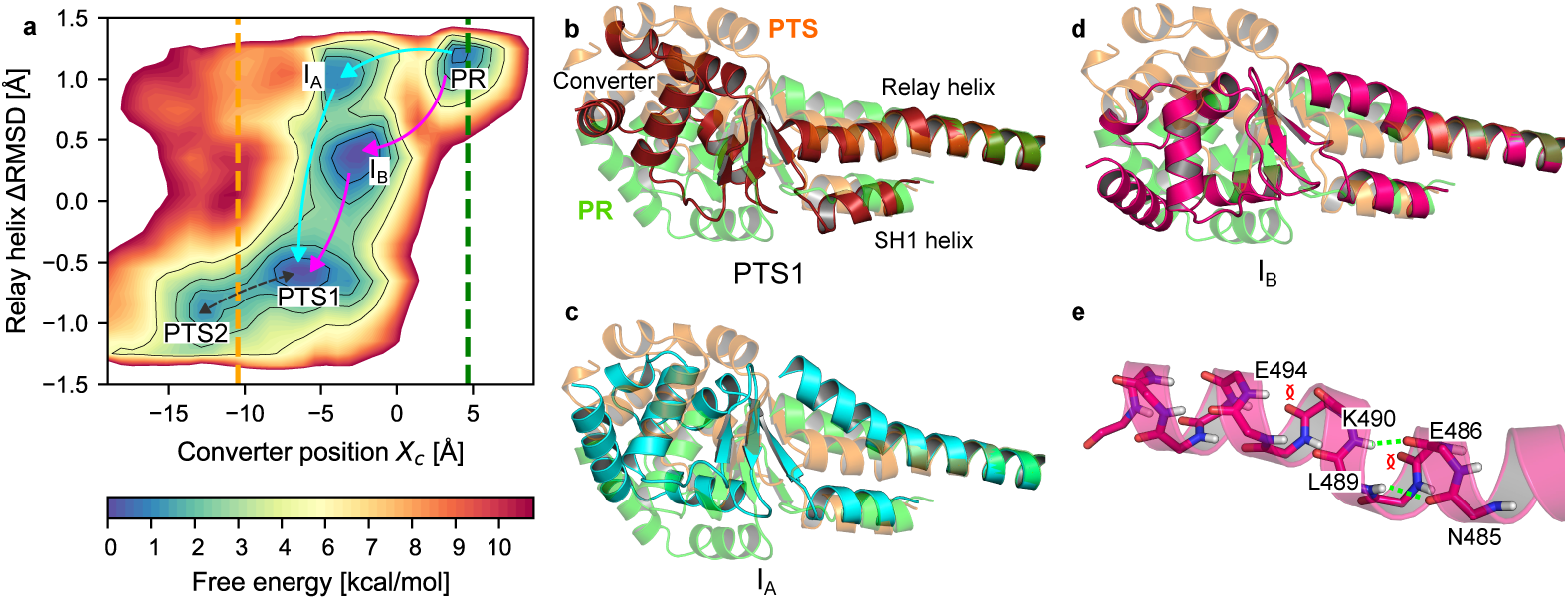
Analysis of the PR → PTS transition by eABF. (a) Free energy landscape obtained by eABF with metastable states labelled. The dotted green (respectively orange) line materializes the value of *X_c_*in the PR crystal structure (respectively PTS crystal structure). The arrows indicate putative transition pathways, namely Path *A_ABF_* (cyan) and Path *B_ABF_* (pink). (b-d). Representative conformations of the converter and Relay and SH1 helices of the PTS1 (b, dark red), I*_A_* (c, cyan) and I*_B_* (d, bright pink) metastable states, compared to PR (transparent green) and PTS (transparent orange) equilibrated structures. Structures are aligned onto the N-terminal fragment of the Relay helix (residues 468-482) to better isolate changes in Relay and SH1 helix orientation and converter position. (e) Close-up on the disruption of the Relay helix in I*_B_*.

### Transition tube and mechanism

The free energy landscape shows that a low free energy transition tube connects PR to PTS. This indicates that the PTS state is kinetically accessible from PR and supports the hypothesis that it is an on-pathway intermediate along the recovery stroke. Two local free energy minima are identified within the transition tube, labelled I*_A_* and I*_B_* on Fig 2a. We use structural clustering to reveal their structural features. This analysis indicates that I*_A_* corresponds to an intermediate along the PR→ PTS transition where the converter has moved by 7.8 Å in the re-priming direction, promoting partial tilting of SH1 and bending of the RH in response, but not yet formation of the kink (Fig 2c). In I*_B_*, the converter has moved by only 6.5 Å and its orientation is quite different from I*_A_*, and closer to PR ((Fig 2d and Supp. Fig. S5a). Moreover, in I*_B_* the RH backbone differs both from the regular helical structure observed in PR and the kink observed in PTS (and PPS) crystal structures. Rather, the RH exhibits local unfolding and bending near residues 490 to 494 (*i.e*, ∼5 residues in C-ter of the canonical kink), see Fig 2e and Supp. Fig. S5b.

Although the transition tube is suggestive of a sequential mechanism with I*_A_*, then I*_B_* as on-pathway intermediates, the differences in converter position/orientation and RH structure are also consistent with I*_A_* and I*_B_* belonging to two different pathways (Paths *A_ABF_* and *B_ABF_*, respectively in cyan and pink on Fig. 2). In Path *A_ABF_*, the initiating event is a ≈8 Å movement of the converter which crosses the highest free energy barrier along this path (6 ± 1 kcal mol^−1^, see SI for statistical error estimation), suggesting it may represent the rate-limiting step. The second step to reach PTS1 is the formation of the kink in the RH, which contributes an additional ∼2 Å of converter swing. In Path *B_ABF_*, the first step is a 6.5 Å movement of the converter associated with local disruption of the RH backbone, and the second step corresponds to the formation of the ”canonical” kink in the RH along with a further 3.5 Å converter swing to PTS1. Despite their differences, both paths predict an early converter movement as the first step of the recovery stroke in Myo6. Notably, our previous unbiased simulations of the PR state did capture spontaneous converter uncoupling without Relay helix kinking [26], which may indicate that Path *A_ABF_*is more likely to represent the dominant mechanism to reach PTS (Supplementary Figure S4, simulation PR (3)).

The analysis of the free energy landscape allows us to 1) validate the relevance of PTS as an accessible metastable state from PR and 2) draw a first mechanistic picture of this transition. However, it does not reveal how other important structural rearrangements, such as changes in the SH1 helix, are coupled to the transition. We now address this with the string method in collective variables.

### String method analysis of the PR → PTS transition

First, we performed preliminary string optimizations in (*X_c_,* Δ*RMSD*), which showed that the minimum free energy path (MFEP) for the PR → PTS transition lies within the transition tube predicted by eABF (Supp. Fig. S7, Supp. Fig. S8, Supplementary Text 2). This independent agreement confirms that the general features of the transition predicted by eABF are robust. Then, we set out to describe the transition mechanism with higher resolution. To this end, we used the string method with 12 collective variables giving a more detailed insight into the rearrangements involved in the PR → PTS transition (see Table 1; we note that Δ*RMSD* was not used as a supporting CV for 12D CVSM calculations because the 4 kink distances serve the same purpose, but with better descriptive power).

**Table 1.** List of collective variables defined in the study of the myosin VI recovery stroke. All distances, orthogonal projections and RMSD are in Å. All angles are in deg.

| Collective Variable | Rearrangement | Biased in | Force constant<br>(kcal/mol/U<sup>2</sup>) |
| --- | --- | --- | --- |
| Orthogonal projection $X_c$ | Converter swing | eABF, string (2d, 12d, 20d) | 10 |
| Orthogonal projection $Y_c$ | Converter swing | string (12d, 20d) | 10 |
| Orthogonal projection $Z_c$ | Converter swing | string (12d, 20d) | 10 |
| Distance $i_1$ (65O:707OG) | Converter/main body interaction | string (20d) | 20 |
| Distance $i_2$ (66CE:F763 phenyl ring) | Converter/main body interaction | string (20d) | 20 |
| Distance $i_3$ (65O:702NE2) | Converter/main body interaction | string (20d) | 20 |
| Distance $i_4$ (66O:702NE2) | Converter/main body interaction | string (20d) | 20 |
| Distance $i_5$ (64O:761N) | Converter/main body interaction | string (20d) | 20 |
| Orientation angle $\theta_{RH}$ | Orientation of the Relay helix | string (12d, 20d) | 5 |
| Orientation angle $\theta_{SH1}$ | Orientation of the SH1 helix | string (12d, 20d) | 5 |
| Distance $d_{R/SH1}$ (469-482CA:693-703CA) | Relay/SH1 seclusion | string (12d, 20d) | 20 |
| Orthogonal projection $Z_{SH1}$ | SH1 piston motion | None | N.A. |
| Relay helix $\Delta RMSD$ | Relay helix kink | eABF, string (2d) | 125 |
| Distance $k_1$ (486O:490N) | Relay helix kink | string (12d, 20d) | 50 |
| Distance $k_2$ (485O:489N) | Relay helix kink | string (12d, 20d) | 50 |
| Distance $k_3$ (485O:490N) | Relay helix kink | string (12d, 20d) | 50 |
| Distance $k_4$ (486O:491N) | Relay helix kink | string (12d, 20d) | 50 |
| Distance $k_5$ (490O:494N) | Relay helix secondary kink | string (20d) | 50 |
| Distance $k_6$ (491O:495N) | Relay helix secondary kink | string (20d) | 50 |
| Distance $k_7$ (492O:496N) | Relay helix secondary kink | string (20d) | 50 |
| Dihedral $\chi_{11}$ (L489 N:CA:CB:CG) | Hydrophobic switch | string (12d, 20d) | 0.01 |
| Dihedral $\chi_{12}$ (L700 N:CA:CB:CG) | Hydrophobic switch | string (12d, 20d) | 0.01 |
| Distance $d_1$ (R205CZ:E461CD) | Critical salt-bridge | None | N.A. |
| Distance $d_\gamma$ (G459N:ATPO1G) | Switch II-ATP hydrogen bond | None | N.A. |

The starting (or guess) path for string optimizations was obtained by ”uplifting” and regularizing the 2-dimensional string optimized over the eABF free energy landscape to the 12-dimensional space (see Supplementary Text 2). Two separate string optimizations, with slightly different parameters, were performed until convergence (Supp. Fig. S9, Supplementary Text 2). Visual analysis of the structures along averaged strings over the last 50 iterations shows that the two calculations converged towards a similar sequence of events, which represents the most likely pathway for the PR → PTS transition of Myo6, and which we call Path A. Backprojection of the two converged Path A strings (Strings A1 and A2) onto the (*X_c_,* Δ*RMSD*) plane reveals that they both lie within the eABF-predicted transition tube. Interestingly, the two converged strings visit *I_A_*, but not *I_B_*, see Supp. Fig. S10. To obtain an energetic view of the transition mechanism, we then performed an umbrella sampling calculation along string A1 with a finer discretization (from 32 to 128 equally-spaced images) and computed the free energy profile along the string, and 2D free energy maps along pairs of observables (see Methods and Supplementary Text 3). From the analysis of string A1, we now describe the sequence of events, structural couplings, and energetics governing the PR → PTS transition.

### Overview of the mechanism

#### Free energy profile and comparison with eABF

The free energy profile along string A1 computed by Umbrella Integration [39] shows 4 metastable states (Fig. 3a), indicating a 3-stage mechanism. Projection of the umbrella sampling simulations along string A1 onto the (*X_c_,* Δ*RMSD*) with MBAR [40, 41] reveals a free energy landscape very similar to the eABF one, but with a number of remarkable differences (Fig. 3b). Free energy minima corresponding to states PR, I*_A_* and PTS1 are identified. PTS2 is not detected because the string does not extend to this basin. I*_B_* is not seen despite its position in configurational space being clearly sampled. This confirms the proposal that *I_A_* and *I_B_* do not belong to the same transition pathway. Comparison of I*_A_* representative structures sampled independently in the eABF and string calculations reveals striking similarity (Supplementary Figure S6), which confirms that the same metastable intermediate is picked up by both calculations and justifies using the same denomination indifferently. We conclude that Path *A_ABF_*and Path A are virtually identical, and we can now detail the transition mechanism.

**Fig 3.**
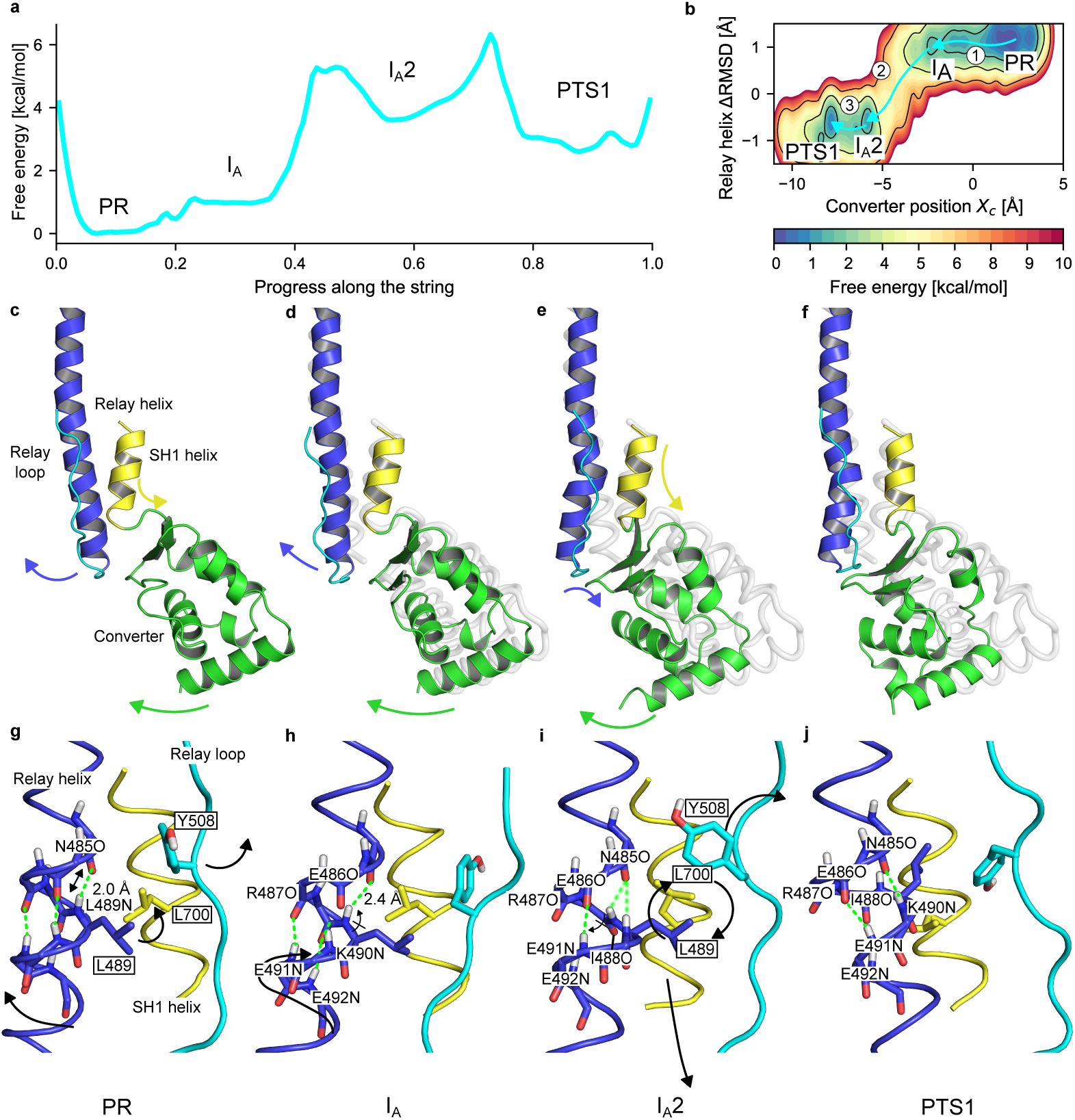
Free energy profiles and mechanism of the PR → PTS transition. (a) PMF along the path obtained by Umbrella Integration. (b) 2D PMF along (*X_c_,* Δ*RMSD*) obtained by MBAR. Arrows indicate transition steps between metastable states. (c-f) Sequence of metastable states in the PR → PTS transition from string calculations. Arrows indicate the movements of structural elements. For easier comparison, the PR structure is shown in transparent grey. (g-j) Mechanism of formation of the kink in the RH. Arrows indicate backbone and side chain movements. H-bonds are shown as green dotted lines, shading indicating weak binding. (c,g) From PR, in response to the first converter movement, the RH C-ter shifts and rotates, which extends the N485O:L489N H-bond and brings L489 closer to L700. (d,h) In I*_A_*, the second converter movement translates into corkscrew movement of the RH along its axis, shifting K490N and E491N by a quarter of helical turn, but complete rotation of RH is prevented by the L489/L700 contact. Instead, the I488-L489 peptide bond flips, which breaks the I488O:E492N H-bond. This deprives R487O and I488O from H-bond partners, and weakens the N485O:L489N H-bond because of unfavorable angle. In counterpart, non-canonical H-bonds E486O:E491N and N485O:K490N form. (e,i) In I*_A_*2, structural tension is relieved by movements of the SH1 helix and the Relay loop. This frees sterical constraints on the L489 and L700 side chains, which exchange their positions while remaining in contact. In addition, I488-L489 flips back, allowing the I488O:E492N H-bond to reform. (f,j) In PTS1, the hydrophobic switch is rearranged, stabilizing the kink in the Relay helix. R487O is free of H-bond partner. previously reported [26]. Although I*_A_* is in fast interconversion with PR, it represents an important intermediate because converter uncoupling from the main body is a pre-requisite to facilitate its swing.

#### Step 1: PR → *I_A_*: converter rotation re-orients Relay and SH1 helices

Starting from the PR state, the first step is a thermally-activated (stochastic) ≈8 Å movement of the converter leading the motor domain into the I*_A_* intermediate (Fig 3c,d, Supp. Fig. S11). This is consistent with the eABF prediction, but the free energy barriers differ, with umbrella sampling indicating a ≈1 kcal mol^−1^ barrier (Fig 3a), much smaller than the 6 kcal mol^−1^ eABF barrier (Fig 2). The converter displacement reduces the total number of contacts it makes with the N-terminal subdomain (Supp. Fig. S12a,b,e) which increases the converter’s positional freedom relative to PR, because it is attached to the main body only through the flexible Relay and SH1 helices. The most energetically costly event in the PR → I*_A_* step seems to be the disruption of the converter/N-terminal contacts. Since these are mostly non-specific hydrophobic interactions, the corresponding free energy barrier is rather small and can probably be crossed reversibly on relatively short timescales. This is consistent with spontaneous ∼100 ns converter uncoupling in unbiased MD of PR we

Both the RH C-ter moiety (residues 489 to 499) and the SH1 helix re-orient concurrently with the converter movement (Fig 3c,d, Supp. Fig. S11). The C-ter moiety of the RH is strongly bound to the converter through specific contacts, and its N-ter moiety is anchored within the main body. As a result, the RH bends following the movement of the converter. RH bending entails only limited backbone perturbation, with a transient extension of the N485O:L489N hydrogen bond close to the bending point (Fig 3g,h, Supplementary Figure S11). This enables a ”corkscrew”-like rotation of the RH C-ter moeity around the helical axis, which brings the L489 side chain in contact with L700 on the SH1 helix, resulting in tilting of the SH1 helix. This steric clash constitutes a first blocking point to the free movement of the converter, which is later relieved through extensive conformational changes.

#### Step 2: *I_A_* → *I_A_*2: converter movement drives formation of the kink

At the beginning of Step 2, a second movement of the converter is translated into further angular re-orientation of RH and SH1 (Fig 3d,e, Supp. Fig. S11). However, unlike Step 1, the RH backbone can no longer accommodate the strain only through bending, because this movement is blocked by the clash of L489/L700 side chains. Therefore, the ”corkscrew”-like motion of the RH backbone is coupled with dramatic alterations to the H-bonding pattern, namely the formation of the E486O:E491N and the disruption of the E486O:K490N, R487O:E491N, I488O:E492N H-bonds (Fig 3h,i). N485O:L489N, which was extended by about 0.5 Å in I*_A_*, returns to perfect geometry. And, atoms N485O and K490N are brought closer but not yet quite enough to form a H-bond. The cost of breaking these backbone H-bonds likely explains why this step would be rate-limiting for the PR → PTS transition, as suggested by the free energy profile (highest free energy barrier, ≈4 kcal mol^−1^, Fig. 3a). Upon reaching *I_A_*2, the Relay helix is clearly kinked, even though the H-bonding pattern differs slightly from what is observed in PTS. Notably, I*_A_*2 features outward-pointing, unbound R487O and I488O backbone atoms, which may explain why I*_A_*2 is rather high in free energy (Figure 3a and Fig 3i), even if the formation of at least one non-canonical H-bond in the RH backbone (E486O:E491N) makes this state metastable.

The analysis of the I*_A_* → I*_A_*2 step indicates that the main driving force for the formation of the RH kink is the movement of the converter. It is plausible that capturing a large enough converter movement to drive the costly kink formation would require first putting the converter in an uncoupled state. There, the converter can efficiently harness fluctuations from the bath and amplify them into functional rearrangements within the motor domain. This shows why it is important to first visit I*_A_* as an easily accessible uncoupled state.

#### Step 3: *I_A_*2 → PTS1: seclusion of the SH1 helix and rearrangement of the hydrophobic switch

The tight packing of the L489/L700 side chains is preserved throughout the transition to I*_A_*2, including upon formation of the kink. This builds up structural frustration at the Relay/SH1 interface. During the I*_A_*2 → PTS1 transition, this frustration is relieved through two consecutive rearrangements. First, the SH1 helix moves by ≈1 Å parallel to the RH in a ”piston-like” motion, and re-orients upward by ≈5 deg (Fig. 3e,f, Supplementary Figure S11). This creates enough space to allow packing L489 against the SH1 helix. Second, a movement of the Relay loop relieves the sterical hindrance from the Y508 side chain (Fig 3i,j). This allows the L489 and L700 side chains to eventually exchange their positions (Fig 3i,j). In the process, the I488-L489 peptide bond rotates, allowing I488O to re-form a H-bond with E492N while the N485O:L489N hydrogen bond breaks. Fischer et al. and Baumketner independently proposed that a similar transition for the Dd Myo2-homologous side chains (Dd Myo2 F487, F506 and I687) stabilizes the kink [14, 24]. By analogy to the ”aromatic switch” of Fischer et al., we call this triplet of side chains the ”hydrophobic switch” for Myo6. The sequence of events captured by our calculations for L489 and L700 is reminiscent of the mechanism proposed by these investigators, but not entirely consistent with it (see Discussion). Because the Relay loop is very flexible, its movement is likely to be a stochastic, low-barrier event weakly coupled to the rest of the transition. Thus, the movement of the SH1 helix, driven by the push of L489, likely accounts for most of the ≈3 kcal mol^−1^ barrier (Fig. 3).

### No coupling to the active site is detected

We searched for a structural response of the active site to the rearrangements of the PR → PTS transition. First, we computed the free energy map along the distances defining the catalytically-competent configuration from umbrella sampling (Fig. 4a,c), which shows that the position of Switch II is nearly unaffected by the transition, like our observations from eABF. Then, we looked into a possible communication pathway between SH1 and the active site proposed by Fischer et al. for Dd Myo2 [17]. These authors proposed that the ”piston-like” motion of the SH1 helix translates into the formation of a hydrogen bond between P-loop and Switch II, which may contribute to ATPase activation (see also Discussion). To assess whether a similar mechanism is at play during the PR → PTS transition, we computed the free energy map along the SH1 helix longitudinal projection *Z_SH_*_1_ (accounting for the piston-like motion) versus the S153N:F460O hydrogen bond distance between P-loop and Switch II, see Fig. 4b,d. Although the free energy map suggests that the piston movement brings S153 and F460 slightly closer, this does not result in the formation of a hydrogen bond between these two residues. The contact between the wedge loop and the SH1-SH2 junction was proposed as the key element mediating structural communication between Switch II and the SH1 helix by Fischer et al. [17]. In our simulations, F582 on the wedge loop is brought somewhat closer to the SH1-SH2 junction as the transition proceeds (Fig. 4e), but this is not sufficient to trigger the global movement of the wedge loop which would drive the formation of the P-loop / Switch II hydrogen bond. We conclude that the near-complete rearrangement of the mechanical amplifier element (Relay/SH1/Converter region), which is required to reprime the lever arm, proceeds without detectable effect on the active site.

**Fig 4.**
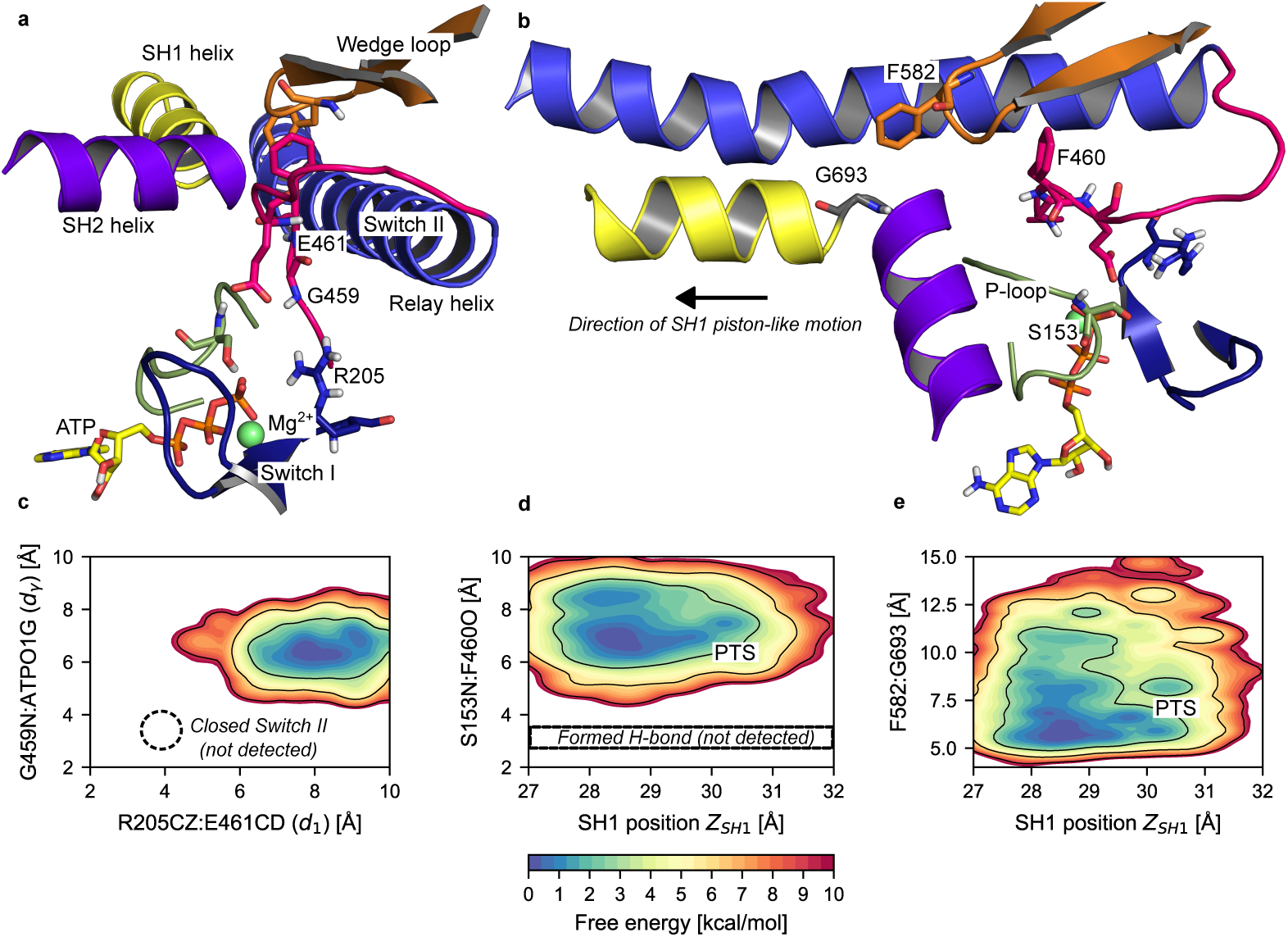
The active site remains open throughout the PR→PTS transition. (a) Close-up on the active site for a representative PTS configuration from umbrella sampling. The critical salt-bridge (R205CZ-E461CD, distance *d*_1_) and the Switch II-ATP hydrogen bond (G459N:ATPO1G, distance *d_γ_*) are clearly not formed, corresponding to an ATPase-incompetent active site. (b) Same configuration seen from a different angle, highlighting the SH1-SH2 junction, the wedge-loop and the P-loop. Although F582 is close to G693 at the SH1-SH2 junction, the P-loop-Switch II hydrogen bond (S153N:F460O) is clearly not formed. (c) 2D PMF over *d*_1_ and *d_γ_* computed along the transition. The closed Switch II state is not detected. (d) 2D PMF over the piston-like movement of the SH1 helix (*Z_SH_*_1_) and the P-loop-Switch II hydrogen bond distance (S153N:F460O) computed along the transition. The formed hydrogen bond is not detected. (e) 2D PMF over the piston-like movement of the SH1 helix (*Z_SH_*_1_) and the distance between the wedge loop and the SH1-SH2 junction (F582-G693 distance). The completion of the piston-like movement of SH1 favors closer approach between the wedge loop and the SH1-SH2 junction.

### Alternate mechanism for the PR → PTS transition

To assess the robustness of our findings to the starting point of string optimizations, we carried out a second set of string calculations where the guess path was obtained as the straight path joining PR to PTS in 12D CV-space (Supplementary Text 2). This resulted in the so-called Path B, which is also consistent with the eABF transition tube, but does not overlap with Path A. Instead, Path B admits I*_B_* as an intermediate and is virtually identical to Path *B_ABF_*, *i.e.,* a mechanism in which the RH transiently develops a secondary kink (see Supplementary Text 4 and Supplementary Figure S14). Because Path B entails more structural perturbation than Path A (two kinks instead of one), it is less likely to represent the main sequence of events of the PR→PTS transition, hence why our focus is on Path A. Path B may contribute marginally to the transition flux, but this does not change our conclusions that lever arm priming comes from harnessing converter fluctuations without immediate coupling to the active site.

## Discussion

Molecular motors like myosin convert chemical energy from ATP hydrolysis into mechanical work following established thermodynamic principles [42]. However, how these principles are implemented at the molecular level remains unclear. We have tackled this question using all-atom MD simulations and free energy calculations. In particular, we focused on the recovery stroke transition that couples the re-priming of the myosin lever arm to ATP hydrolysis. Our results shed light on the chemo-mechanical transduction mechanism and illuminate structural intermediates visited during this off-actin transition, which may be targeted by allosteric modulatory ligands [43, 44].

Using advanced free energy simulations started from the X-ray structure of the post-rigor (PR) state of myosin VI, we illuminated a low-energy transition pathway connecting PR to a previouly characterized structural intermediate, named PTS, with an unprecedented level of detail. Consistent with previous observations [26], our calculations suggest that the recovery stroke of myosin VI is initiated by a movement of the converter by thermal fluctuations accompanied by the kinking of the Relay helix, which corresponds to the highest energy barrier, tilting of the SH1 helix and stabilization via a switch of hydrophobic interactions between residues at the Relay/SH1 interface. Remarkably, these calculations not only ”re-discover” PTS agnostically but also show that the same sequence of events could be obtained using two different free energy simulation methods, which strenghtens the relevance of these findings. Most importantly, the striking observation is that repriming of the converter subdomain, *i.e.,* a key element of the mechanical amplifying region of the motor protein, precedes closure of Switch II at the nucleotide-binding site. In addition, our free energy analysis predicts that the rate-limiting step to PTS is ≤6 kcal mol^−1^ with Switch II open, whereas the free energy barrier for closing Switch II from PR was 12 kcal mol^−1^ [26]. Assuming similar pre-exponential factors, these results suggest that the converter-initiated mechanism would be ≈23 000 times faster than the Switch II-initiated scenario. Intriguingly, these observations suggest that lever-arm repriming and ATP hydrolysis are only weakly coupled, so that the recovery stroke transition can be initiated by thermal fluctuations and driven via modulation of kinetic barriers. These observations thus support the conclusion that the recovery stroke in Myo6 is mediated by a ”ratchet-like” mechanism. As compared to a Switch II-initiated mechanism, a ratchet-like mechanism would increase the motor efficiency by ensuring that ATP hydrolysis takes place only once the mechanical amplifier region is re-primed,+ avoiding futile ATP consumption [45], and allowing the motor to effectively harness and ”rectify” ubiquitous Brownian fluctuations.

Our mechanistic proposal contrasts with previous models of the recovery stroke, most of whom predicted Switch II closure as the initiating event. The mechanism proposed by Fischer and co-workers in Dd Myo2 consists of two phases [14, 17]. In phase I, a movement of the converter couples to Switch II closure through a rigid-body ”seesaw” movement of the RH. In phase II, an additional movement of the converter couples to SH1 helix tilting, RH kink formation, and the establishment of Switch II/P-loop interactions. This scenario is plausible and consistent with Dd Myo2 mutants, but it derives from zero-temperature calculations, which are inherently biased towards strong coupling. Elber and West calculated a zero-temperature transition path similar to Fischer’s model and used finite-temperature MD to evaluate the transition rate along this path [22]. Although the predicted rate is consistent with experimental estimates, these simulations are likely too short (a few ps) to appreciably deviate from the zero-temperature transition path. Woo and Harris explored the recovery stroke of Scallop myosin II by umbrella sampling. These calculations suggested a weak coupling between Switch II closure and converter rotation, but it is unclear whether the short simulation trajectories (*<*6 ns) could faithfully reveal the sequence of events and the energetics [18, 19]. Using Targeted MD (TMD) simulations from the PR structure of Dd Myo2, Cui and co-workers predicted an early movement of the converter followed by RH kinking and Switch II closure [21], which is reminiscent of the mechanism proposed here. However, partial Switch II closure (critical salt-bridge formation) was observed at the beginning of the transition consistent with an early Switch II closure. Baumketner and Nemeslov came to the same conclusion using high-temperature simulations of the PR state in implicit solvent [23]. Later, using enhanced sampling Baumketner convincingly showed that rearrangements within the mechanical amplifier region are statistically coupled, but its fragmental model lacking Switch II prevented him from exploring the consequences at the nucleotide binding site [24, 25]. In brief, methodological and sampling limitations (with respect to current state-of-art) may explain why earlier studies did not detect PTS. In addition, the usage of different myosin isoforms might also contribute to diverging outcomes.

The ratchet-like model presented here shares features with the more broadly accepted Switch II-initiated mechanism by Fischer’s and Baumketner’s although with important differences. First, in the Switch II-initiated scenario, the seesaw motion of the Relay helix precedes its kinking, as the latter involves threading the bulky Dd Myo2:F487 between the RH backbone and the Relay loop, which would be sterically prevented until the RH seesaw motion opens the way. By contrast, we find that a movement of the Relay loop is sufficient to enable threading of the homologous L489 in Myo6 before the RH seesaw motion, thus resulting in an opposite order of events. Second, in the Switch II-initiated scenario it was found that SH1 tilting is coupled to the formation of a H-bond between Switch II and P-loop at the nucleotide-binding site (DdMyo2:S183N-F458O) via the wedge loop. Yet, we do not observe the formation of the corresponding H-bond in Myo6 (Myo6:S153N-F460O) and the PTS crystal structure does not exhibit it either, suggesting that it forms during the PTS → PPS transition instead. Third, in the Switch II-initiated scenario, the tilting of the SH1 helix drives the converter rotation, whereas we come to the opposite conclusion. Fourth, in the converter-initiated scenario, the seesaw motion of the Relay helix (coupled to an additional converter movement and Switch II closure) occurs in the PTS → PPS transition, which will be detailed elsewhere.

Based on these observations, the Switch II- and converter-initiated mechanisms emerge as plausible alternative scenarios depending on whether RH kinking occurs before (converter-initiated) or after (Switch II-initiated) the seesaw motion (Fig. 5). In this interpretation, the PTS state would be naturally described as the post-kink, pre-seesaw intermediate along the converter-initiated pathway. A different intermediate with a partially re-primed converter, an unkinked but ”seesawed” Relay helix, and a closed Switch II should be observed along the Switch II-initiated pathway [17]. To the best of our knowledge, the latter has not been observed experimentally yet. Whether the PTS state and ratchet-like recovery stroke are common to all myosins remains an open question. Dd Myo2 mutants interpreted as supporting the Switch II-initiated recovery stroke do not disprove the ratchet-like pathway [26]. The same is true for pathogenic mutations disrupting the converter-Relay interactions in Human `β`-cardiac myosin, which would perturb the recovery stroke regardless of its mechanism, as supported by recent free energy calculations in Dd Myo2 [46] and `β`-cardiac myosin [47]. In addition, recent free energy calculations on `β`-cardiac myosin also identified a PTS-like intermediate [44]. Last, the ratchet-like mechanism is consistent with the structural intermediate of smooth muscle myosin II trapped off-actin by the CK-571 drug [43], which exhibits a partially re-primed converter, an “un-kinked” Relay helix and an open Switch II. To advance on this question, detailed computational investigations of the recovery stroke mechanism in other myosin isoforms [48] are needed. As a bonus, these analyses would enable the characterization of cryptic binding pockets for the design of myosin’s allosteric modulators [44, 49, 50].

**Fig 5.**
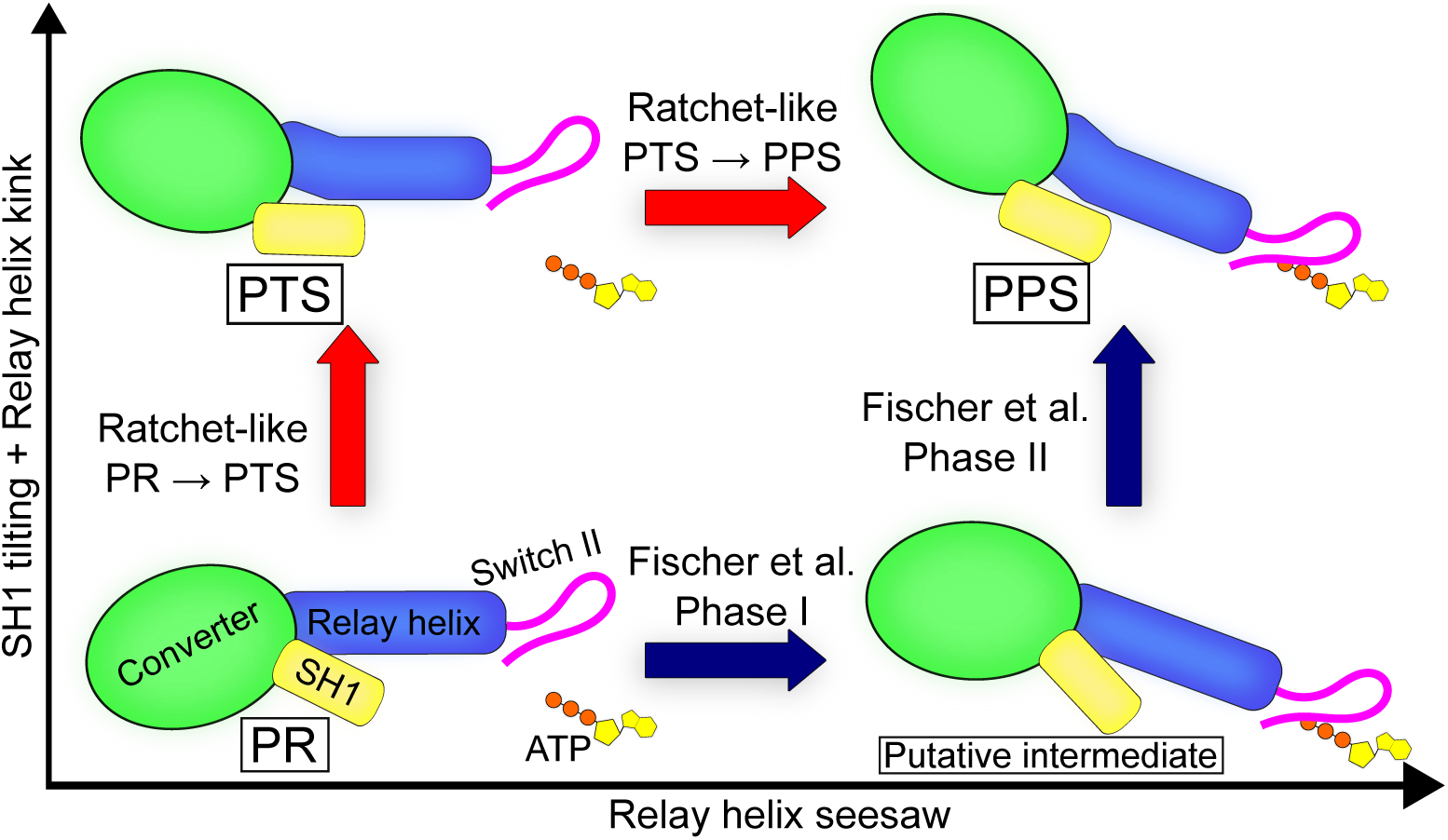
Unified mechanistic framework for the myosin recovery stroke. by a 1 ns-long heating MD simulation to 300 K, then a 2 ns-long equilibration MD at constant temperature (300 K) and pressure (1 bar) during which harmonic restraints were smoothly scaled down. In the present work, only the PR system was used to launch new simulations. Simulations used NAMD 2.13 [51] or Gromacs 2021.5 [52].

## Materials and methods

### Preparation of structural models for Molecular Dynamics simulations

Structural models of the Myo6 PR (PDB ID: 2VAS, [8]) and PTS (PDB ID: 5O2L) states were prepared from the corresponding crystal structures according to the protocol reported in [26]. Briefly, missing loops were built with MODELLER, then each structure was placed in an orthorhombic box of TIP3P water molecules supplemented with Na^+^ and Cl^−^ ions to reach a 150 mM salt concentration. Each system was energy-minimized with harmonic restraints on the protein atoms, followed

### Collective variables

Collective variables (CVs) were used to characterize, and in some cases bias, the structural dynamics of Myo6, focusing on the structural elements involved in the recovery stroke (Fig 1b-e). The complete list of CVs is given in Table 1. All CVs were computed with the *colvars* module [53].

### Converter

The Cartesian coordinates of the converter’s center of geometry in the frame formed by the principal axes of the motor domain are denoted as *X_c_*, *Y_c_*, *Z_c_* (*X*^′^*, Y* ^′^ *Z*^′^ in [26]). They capture the movements of the converter relative to the motor domain during the recovery stroke. The reader is referred to reference [26], SI Appendix, for the full description of the converter CVs. The interactions of the converter with the main body of the motor domain are described by a set of interatomic distances *i_j_* (see Table 1 for details).

### Relay/SH1 elements

Angular descriptors of the orientation of the Relay helix C-terminal fragment *θ_RH_* and the SH1 helix *θ_SH_*_1_ with respect to the PR crystal structure were defined as described in [26], SI Appendix. Following Baumketner [24, 25] we introduced the distance *d_R/SH_*_1_ between the centers of geometry of the CA atoms of residues 469-482 (N-terminal fragment of the Relay helix) and 693-703 (SH1 helix). To characterize the ”piston”-like motion of SH1 highlighted by Fischer and co-workers, we defined *Z_SH_*_1_ as the orthogonal projection of the SH1 helix’ center of geometry onto the longitudinal principal axis of the motor domain (*i.e.*, the same axis as for *Z_c_*). The local conformation of the Relay helix backbone was described by a Δ*RMSD* CV defined as:

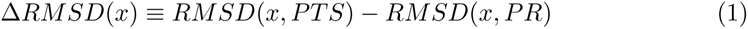

where the *RMSD* values are computed after optimal translation/rotation and with respect to the corresponding crystal structures. For the kink of the Relay helix, Δ*RMSD* was computed on atoms C, CA, O, N (backbone heavy atoms) of residues 485 to 493. This CV takes values ≈1.4 Å in PR (straight Relay helix) and ≈−1.4 Å in PTS/PPS (kinked Relay helix), Fig 1e. Although Δ*RMSD* is defined using the PTS crystal structure as a reference, we note that the local backbone configuration of the kinked Relay helix is virtually indistinguishable between PTS and PPS (0.32 Å RMSD). For all intents and purposes, using a CV based on the PPS crystal structure [7] would have given identical results. To analyze the backbone hydrogen-bonding pattern of the Relay helix with a higher resolution than offered by Δ*RMSD*, we introduced a set of distances *k_j_*(see Table 1 for details). And, we computed dihedral angles *χ*_11_ and *χ*_12_ (see Table 1) to describe side-chain rotameric transitions associated with the formation of the Relay helix kink, as pointed out previously by Fischer et al. and independently by Baumketner [14, 24].

### ATPase site

The opening state of the Switch II loop in the active site was characterized by distances *d*_1_ and *d_γ_* as defined previously [26]. *d*_1_ describes the critical salt-bridge between Switch I (R205) and Switch II (E461). *d_γ_*describes the Switch II-ATP hydrogen bond. When both interactions are formed, Switch II is closed and the motor is considered ATPase-competent.

### Extended Adaptive Biasing Force (eABF) calculations

Two-dimensional eABF calculations were run along *X_c_* × Δ*RMSD* using NAMD 2.13 with the *colvars* module. We prioritized *X_c_*for the eABF calculation as it is the converter component which exhibits the largest variation from PR to PTS, with typical values of 5 Å in PR and −10 Å in PTS/PPS. Briefly, ABF records the average generalized force applied by the system onto the collective variables, then applies an exactly opposite bias to enhance the sampling by flattening the free energy landscape. The free energy profile is recovered by numerical integration of the free energy gradient estimate, which is the opposite of the collected average force. In extended ABF, an extended degree of freedom is harmonically coupled to the collective variable [38]. The generalized thermodynamic force felt by the extended degree of freedom is the harmonic coupling force, which can be easily evaluated. eABF then applies the standard ABF dynamics on the extended degree of freedom. eABF does not require collective variable component orthogonality, which allows us to use a Δ*RMSD* CV. Moreover, it favors smooth convergence of the biasing force estimate [38]. We used the corrected-*z*-averaged restraint (CZAR) estimator to deconvolve the free energy gradient estimate [38] and the *abf_integrate* tool to integrate it. eABF calculations were run with SHAKE active and a 2 fs timestep. The *X_c_* collective variable was coupled to its extended degree of freedom using a 10 kcal*/*mol*/*Å^2^ harmonic force constant; a 125 kcal*/*mol*/*Å^2^ was used for Δ*RMSD*. Force constants were determined by trial-and-error as the lowest force constant sufficient to elicit the corresponding conformational change in short Steered Molecular Dynamics simulations (data not shown). This ensures a coupling that is both strong enough to bias the dynamics, and soft enough to promote smooth exploration of the free energy surface [38]. Both extended degrees of freedom were thermostatted to 300 K with a Langevin thermostat with a 10 ps^−1^ friction constant. The *X_c_* × Δ*RMSD* configurational space was discretized into a 0.75 Å×0.1 Å grid for the calculation. The *fullSamples* parameter was set to 2000 to limit non-equilibrium effects [54].

Similar to our previous work [26, 55], a two-step strategy was used to promote convergence of the eABF calculations. First, a 700 ns exploratory eABF run was performed starting from the equilibrated PR model, yielding near complete sampling of the configurational space. Second, the configurational space was divided into 12 non-overlapping windows separated by harmonic walls. Starting configurations for each window were extracted from the previous 700 ns run; each window was then simulated with eABF (starting with the gradient estimate obtained in the exploratory run) for 500 ns for a total of 6.7 µs of eABF simulation. This improves the sampling and refines the gradient estimate. Convergence was assessed as described in SI.

### String Method in Collective Variables (CVSM) calculations

String method optimizations [31] with the ”swarms-of-trajectories” variant [32] were used to compute MFEPs connecting the PR and PTS states. Strings were defined in either a 2-,12-, or 20-dimensional CV space designed to capture the essential features of the transition while remaining much simpler than full Cartesian space. The supporting CVs are listed in Table 1 along with their harmonic coupling force constant. The CVSM relaxes a discretized path in CV-space following the average drift computed from swarms of short MD simulations; up to unimportant dynamical effects, this average drift occurs along the direction of highest free energy gradient [32, 56, 57]. After each move, the string is reparametrized by enforcing equal spacing of the images, which projects out image displacement colinear to the string and ensures that string evolution takes place only along the orthogonal direction of the free energy gradient. Convergence is reached when orthogonal displacement is zero, that is, when the only image displacement is colinear to the string and thus, exactly cancelled by reparametrization. The converged string then represents the MFEP between the end-states in the supporting CV-space [31]. String calculations were performed using the Tcl scripting interface in NAMD 2.13 and the *colvars* module. Each CV was normalized between 0 and 1 by its total variation to ensure balanced reparametrization. Additional details on the string optimization procedure, parameters, and convergence analyses are reported in SI. Averaged, reparametrized strings over the last 50 iterations were considered for analysis.

### Umbrella Sampling along the converged string

To gain insight into the free energy changes along the converged string A1, we performed Umbrella sampling simulations. First, we used *B*-spline interpolation to increase the resolution of the converged string from 32 to 128 equally-spaced images in normalized CV-space. Then, each image was simulated for 120 ns under harmonic restraints applied to the same set of CVs and force constants as used for string optimization, except that *χ*_11_ and *χ*_12_ were excluded from the set of supporting CVs. Umbrella sampling simulations were run with Gromacs 2021.5 patched with *colvars* [52]. The PMF along the string was computed using Umbrella Integration (UI) [39] with the chain rule as described in [55], see SI. 1- and 2-dimensional PMFs along arbitrary CVs were computed using the Multistate Bennett Acceptance Ratio (MBAR) method [40] with Gaussian kernel density estimation, as implemented in *pymbar* version 4.0 [41].

### Structural clustering

Structural clustering was performed using agglomerative hierarchical clustering with scikit-learn [58]. We used the RMSD over the CA atoms of the Relay and SH1 helices (after optimal translational and rotational fit) as the distance metric. Numbers of clusters were chosen to ensure that at least 2 or 3 clusters with appreciable occupancy (*>*10 %) were obtained.

## Acknowledgments

This project was primarily supported by PRACE allocation ra4332, which allowed priority access to the Curie and Irene supercomputers. Additionally, this work was granted access to the High Performance Computing (HPC) resources of Centre de Calcul Recherche et Technologie (CCRT)/Centre Informatique National de l’Enseignement Supérieur (CINES) under Allocation 2017-[076644] made by Grand Equipement National de Calcul Intensif, and also benefited from computational resources provided by the Mésocentre de Calcul de l’Université de Strasbourg and the Myria supercomputer of the Centre Régional Informatique et d’Applications Numériques de Normandie (CRIANN). The A.H. and M.C. teams were jointly supported by the Fondation pour la Recherche Médicale (Grant DBI20141231319). M.C. was supported by the Agence Nationale de la Recherche (ANR) through the LabEx Chemistry of Complex Systems (Project CSC-MCE-13), and the International Center for Frontier Research in Chemistry. A.H. was supported by a grant from the Association Française Contre les Myopathies 17235. The A.H. team is part of LabEx CelTisPhyBio:11-LBX-0038, which is part of the Initiative d’Excellence Paris Sciences et Lettres (Grant ANR-10-IDEX-0001-02 PSL).

## Supplementary Information

### Supplementary Text 1: Convergence and error analysis for eABF calculations

As seen on Supplementary Figure S1, all relevant regions of configurational space are visited multiple times by the eABF simulations. Convergence of eABF calculations was assessed by monitoring the per-window RMSD of the free energy gradient estimate (Supplementary Figure S2). Most windows representative of the transition tube (*W_11_, W_21_, W_22_, W_32_, W_33_, W_43_*) show a plateau close to 0.5 kcal mol^−1^ Å^−1^ RMSD over the last few tens of ns, suggesting that the generalized force estimate has converged. Bootstrap-like statistical error analysis was performed as in our previous work [26, 59] and shows that statistical error on the free energy values is small (*<* 1.0 kcal mol^-1^ ) everywhere in the relevant regions of 2D free energy landscape, and does not go above 2.0 kcal mol^−1^ even at the irrelevant margins (Supplementary Figure S3). The statistical error at the first free energy barrier is of order 1.0 kcal mol^−1^.

### Supplementary Text 2: String method calculations

#### Procedure

An iteration of the string method with swarms of trajectories consists of the following operations, performed for each image:

1. Equilibration of length *t_eq_* of the system around its image along the string in CV-space using harmonic restraining potentials.
2. Parallel, unrestrained simulation of *n_swarm_*replicas for *t_free_*initiated from the equilibrated system

**Fig S1.**
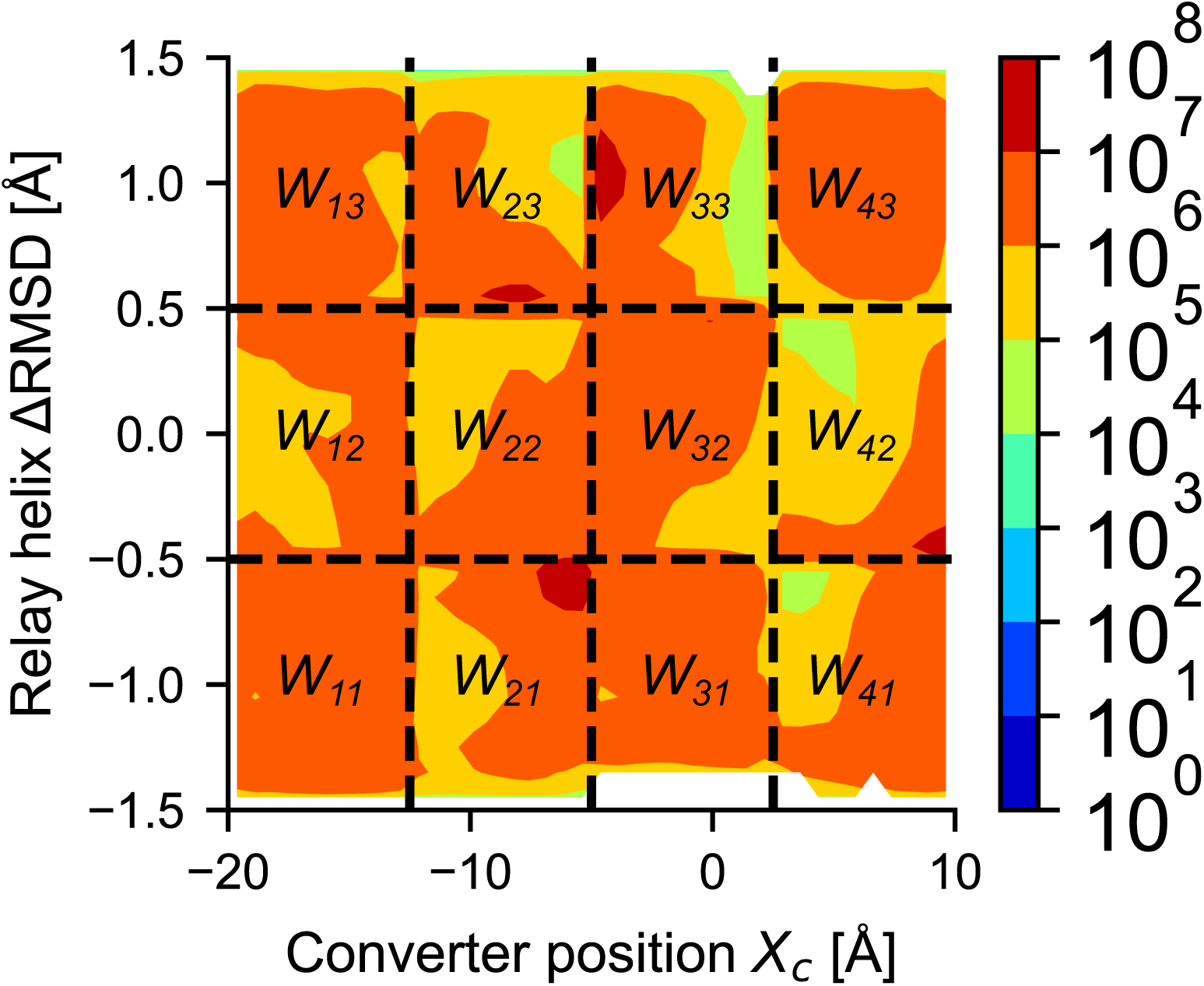
Sampling of the configurational space in eABF simulations. The number of counts, *i.e.*, the number of times a given grid point has been visited, is shown in logarithmic scale. Black dotted lines indicate the window boundaries in stratified simulation. Window labels are shown.

**Fig S2.**
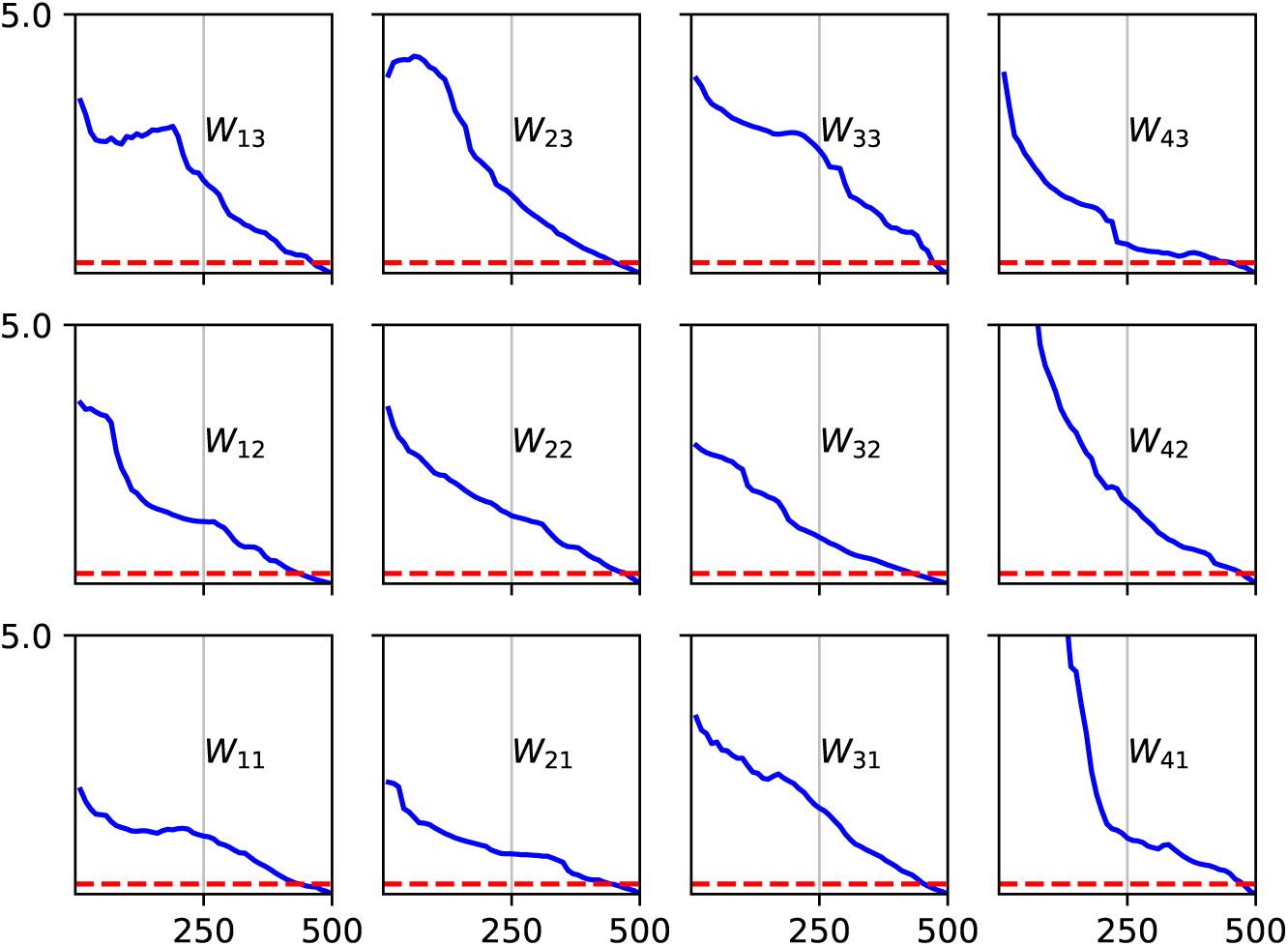
Convergence of generalized force estimate in eABF simulations. Shown is the RMSD of the gradient estimate for each window during the stratified run. X-axis, time of stratified simulation in ns; Y-axis, gradient RMSD, in kcal mol^−1^ Å^−1^. The red dotted lines show the 0.5 kcal mol^−1^ Å^−1^ cut-off which is used to evaluate convergence.

**Fig S3.**
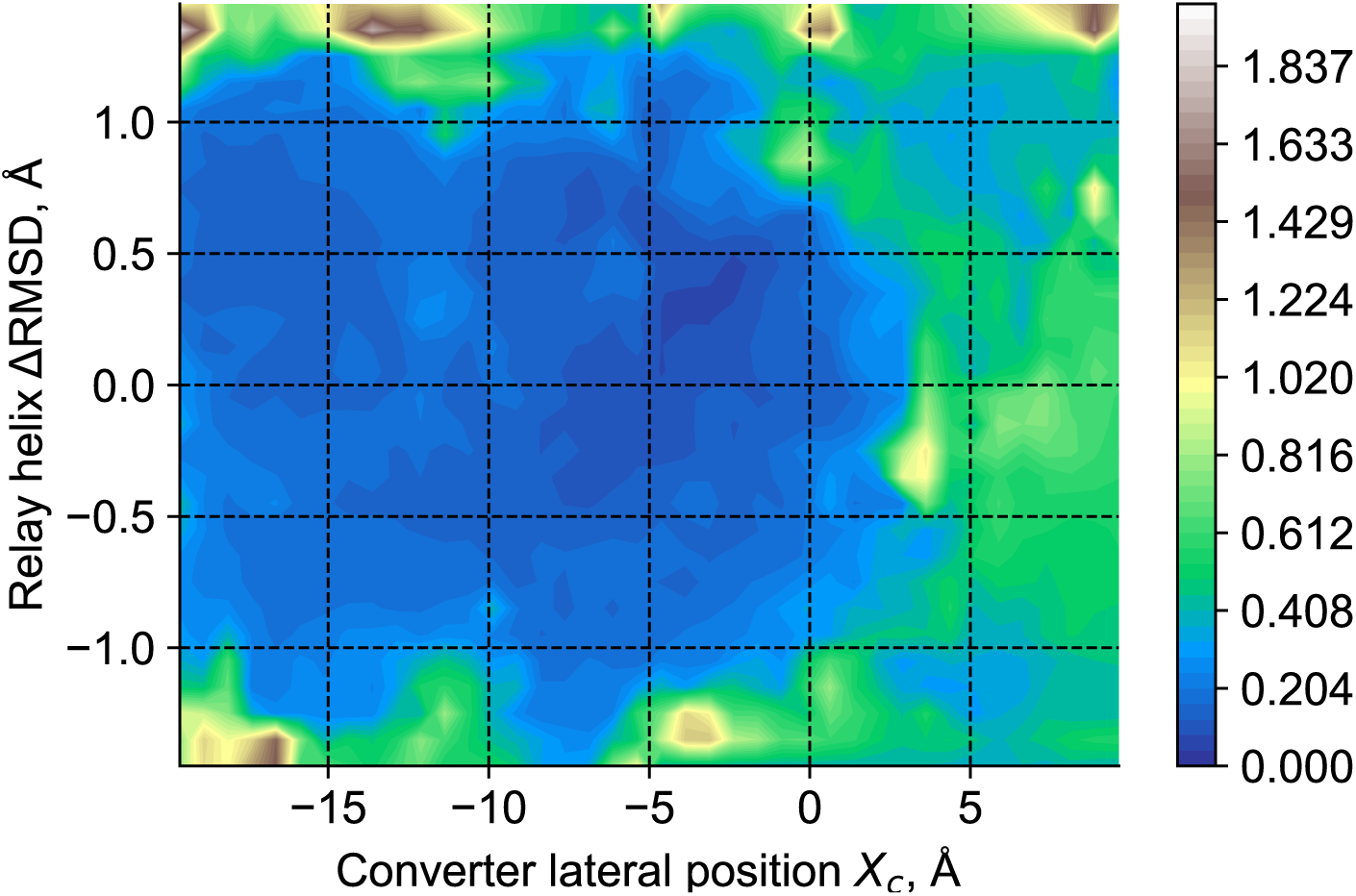
Statistical error estimate for the free energy landscape from bootstrap-like analysis (in kcal mol^−1^).

**Fig S4.**
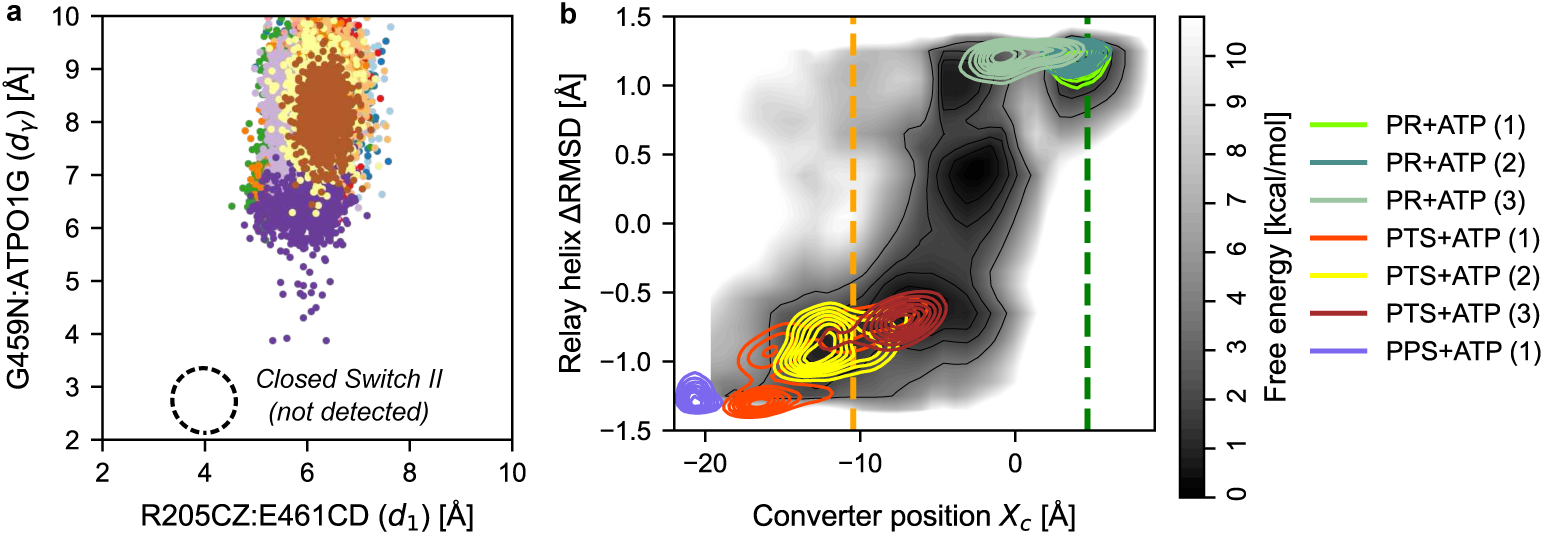
eABF calculations sample open Switch II conformers consistent with independent PTS simulations. (a) Distances *d*_1_ and *d_γ_* never sample the closed Switch II region in stratified eABF simulations. Shown are *d*_1_, *d_γ_* scatter plots colored per window. (b) The PTS basin identified in eABF calculation is consistent with independent, unbiased MD simulations of Myo6. Shown are density lines for *X_c_*and Δ*RMSD* from the Myo6 simulations we reported in [26]. Of note are: the remarkable agreement of the PTS unbiased simulations with the free energy landscape; the sampling of a converter movement in simulation PR+ATP (3) which is roughly consistent with metastable state I*_A_*; and the clear separation of PTS from PPS.

**Fig S5.**
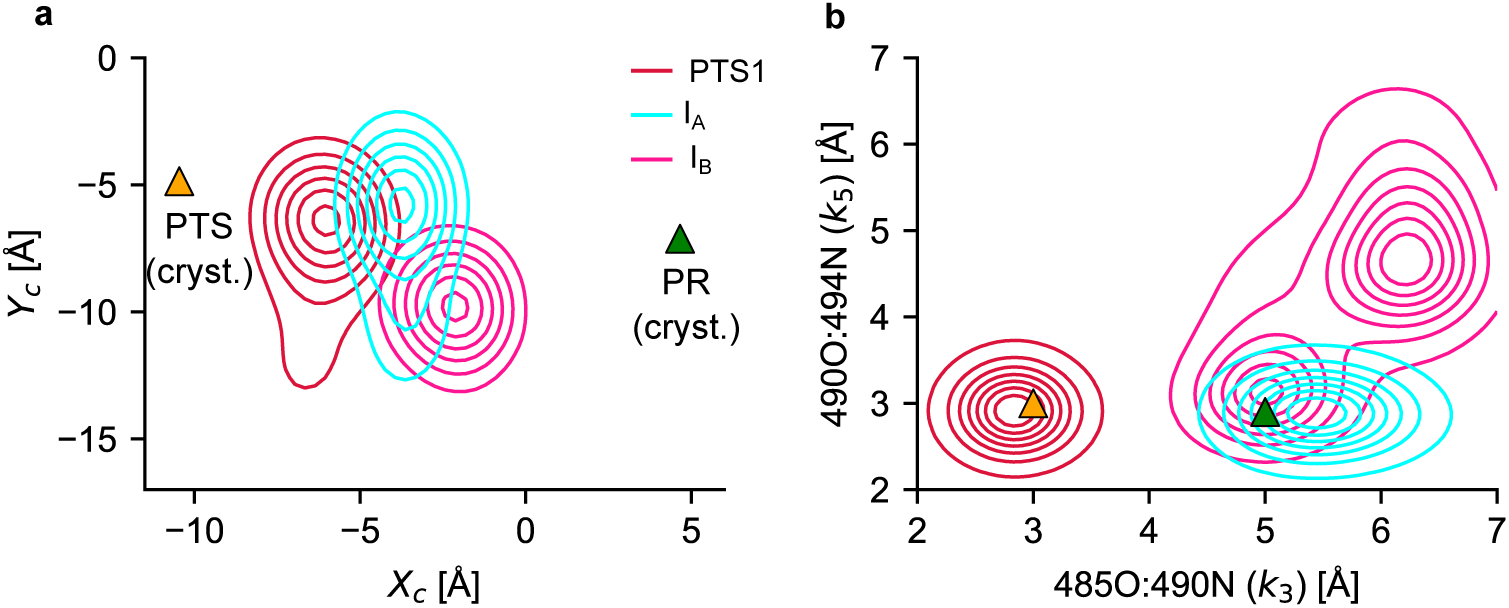
Statistical distribution of selected observables in substates identified in eABF calculations. (a) Distribution of converter coordinates *X_c_* and *Y_c_*. (b) Distribution of Relay helix backbone hydrogen bonds *k*_3_ and *k*_5_. The difference in converter position between I*_A_* and I*_B_*, along with the atypical hydrogen bonding pattern of the Relay helix in I*_B_*, are consistent with I*_A_* and I*_B_* not belonging to the same transition pathway.

**Fig S6.**
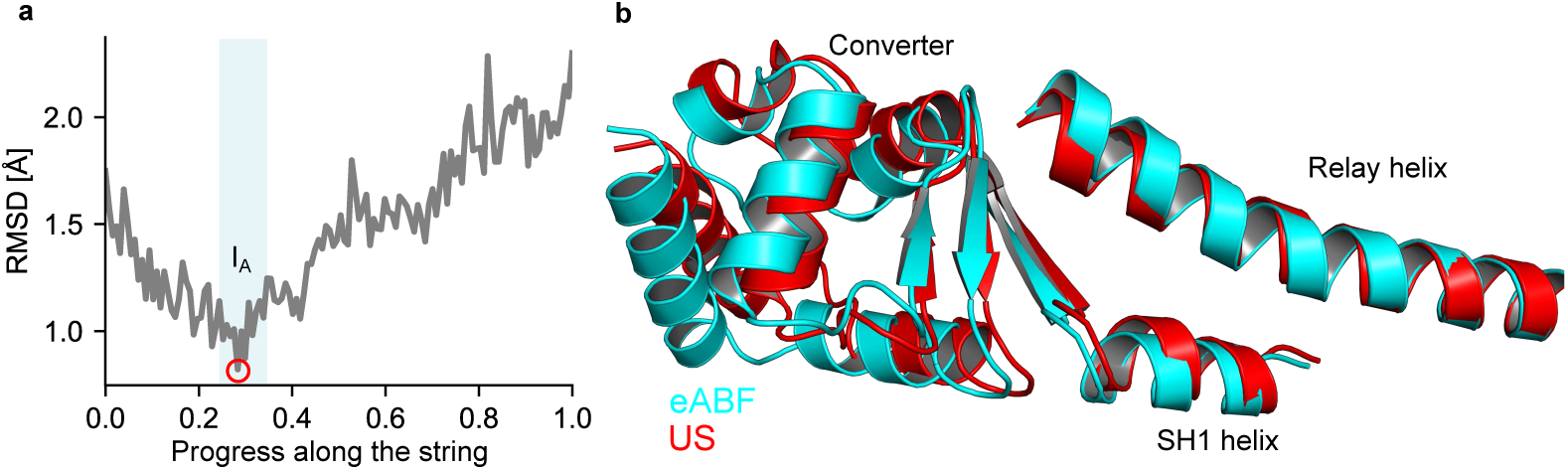
The same I*_A_* intermediate is captured in eABF and string/US calculations. (a) RMSD of structures sampled along String A1 with respect to the representative structure of I*_A_* sampled in eABF, computed after optimal fit on the CA atoms of the Relay and SH1 helices. The RMSD clearly is minimum and below 1 Å when the progress along the path is roughly between 0.24 and 0.35, which corresponds to the I*_A_* basin as identified in string/US calculations. The red dot marks the minimal RMSD value, achieved at image 36 along the string. (b) Structural comparison of the Relay helix, SH1 helix and converter between the representative structure of state I*_A_* sampled in eABF (cyan) and in string/US (image 36, red). The conformation and orientation of the structural elements are extremely close. These results demonstrate that the same metastable basin is sampled in both independent calculations, which justifies referring to them as I*_A_* regardless of their provenance.

For each image, the average drift in CV-space measured over the swarm of unrestrained trajectories is used to update the string. Then, the string is smoothed by local averaging, and reparametrized [31, 32]. This process is iterated until convergence is reached. Convergence of string optimizations is assessed by monitoring the Root Mean Square Deviation (RMSD) of the string in normalized CV-space with respect to the initial string. The string is considered converged when this value stabilizes. String optimization is akin to a gradient-based high-dimensional optimization, for which a well-known issue is the existence of multiple locally optimal solutions. A careful choice of the initial ”guess” path from which string iterations are initialized mitigates this issue by making it more likely to converge to a biologically meaningful solution. Comparing converged strings from different guess path initializations also allows one to assess the robustness of the findings. For these reasons, we consider several different guess paths in our string calculations.

#### String optimizations in **(*X*_c_, Δ*RMSD*)** space are consistent with eABF results

To validate the transition tube predicted by eABF, we first performed string optimizations in (*X_c_,* Δ*RMSD*)-space.

##### Stability of the eABF-predicted MFEP under explicit CVSM dynamics

First, we sought to assess whether the MFEP computed analytically over the eABF free energy landscape were stable under explicit (*i.e.*, atomistic) CVSM dynamics. We constructed this path (hereafter eABF-MFEP) by 1) evaluating analytical derivatives of the eABF free energy landscape and 2) relaxing a 2D string on this free energy landscape using the zero-temperature string method [60] in normalized coordinates. These operations were implemented in a Python/Scipy script [61, 62]. Then, we prepared equilibrated Myo6 conformers along this path in (*X_c_,* Δ*RMSD*)-space as follows. We ”pulled” Myo6 (from the equilibrated PR model) along the eABF-predicted MFEP using a 120 ns-long Steered MD simulation on *X_c_* and Δ*RMSD* with the harmonic force constants reported in Table 1. 32 conformers equally-spaced in normalized CV-space were extracted from the Steered MD trajectory and further equilibrated by 20 ns of umbrella sampling with the same force constants. The resulting conformers were then used to run 2 separate CVSM calculations in 2D space (Supplementary Table S1). In both cases, the path obtained by averaging the strings over the last 50 iterations is remarkably close to the eABF-predicted MFEP. Therefore, up to equilibrium fluctuations, the 2D transition path predicted by eABF calculations is essentially invariant under the string method dynamics, independently confirming that it lies close to the MFEP (Supplementary Figures S7, S8).

##### Relaxation of the straight guess path towards the eABF-predicted MFEP under explicit CVSM dynamics

Using a similar protocol, we prepared equilibrated Myo6 conformers equally-spaced along the straight path joining PR and PTS in 2D CV-space. Relaxation of this straight guess path by two independent CVSM runs shows that it consistently evolves towards the eABF-predicted transition tube (Supplementary Figure S7, Supplementary Table S1). We conclude that independent string optimizations in 2D space confirm the important features of the transition mechanism predicted by eABF.

**Fig S7.**
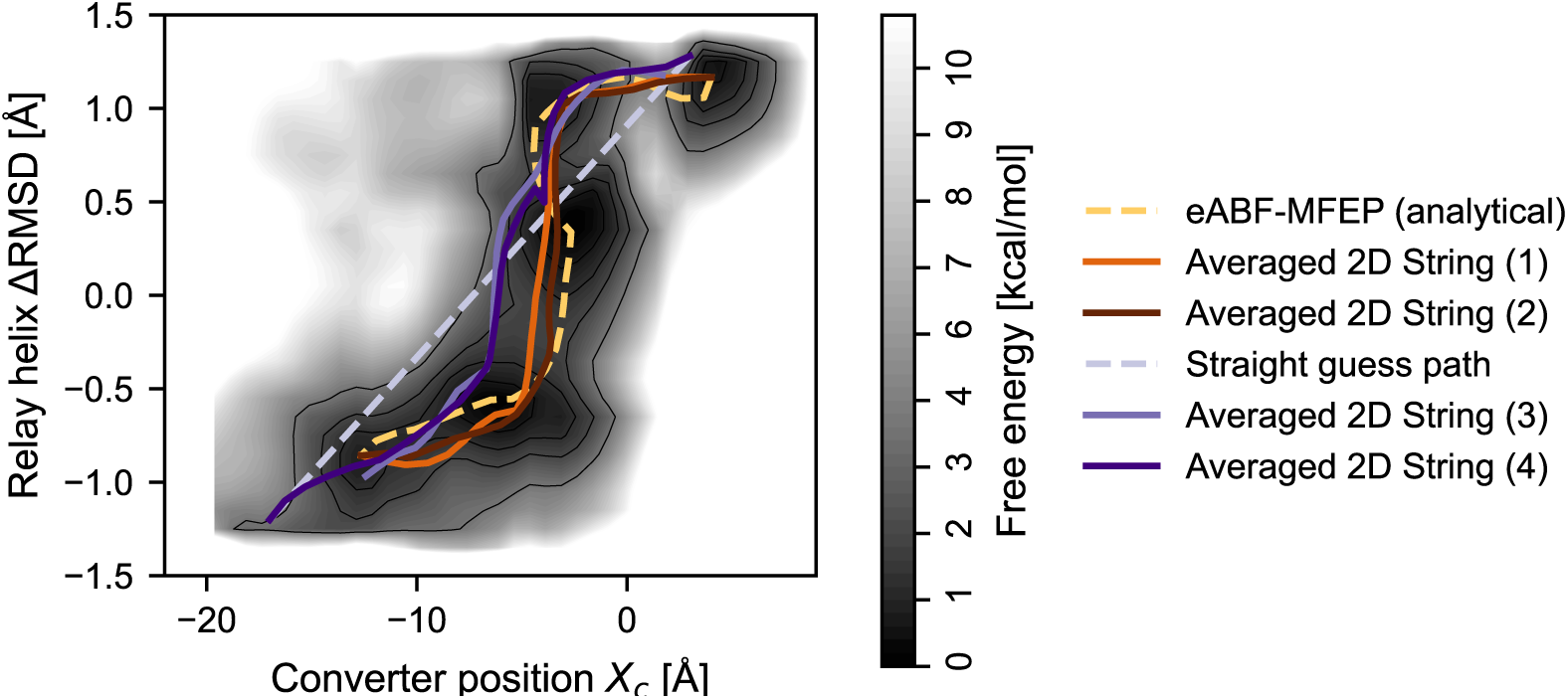
CVSM calculations in 2D CV-space are consistent with the eABF free energy landscape.

**Table S1.** CVSM simulations in 2D CV space.

| Calculation | Guess path | Ends | $t_{eq}$ (ps) | $n_{swarm}$ | $t_{free}$ (ps) | $n_{iter}$ |
| --- | --- | --- | --- | --- | --- | --- |
| 2D String (1) | eABF-MFEP | Fixed | 1 | 10 | 0.5 | 201 |
| 2D String (2) | eABF-MFEP | Fixed | 1 | 10 | 0.5 | 116 |
| 2D String (3) | Straight | Free | 1 | 10 | 0.5 | 127 |
| 2D String (4) | Straight | Fixed | 1 | 10 | 1 | 119 |

#### String method calculations in 12D normalized CV-space

##### Generation of a 12-dimensional guess path from the 2D eABF-MFEP by uplifting

The eABF-MFEP is representative of the general features of the PR → PTS transition, but the 2D CV-space is too coarse-grained to capture the finer details of the mechanism. The 2D MFEP and surrounding transition tube correspond to projections of a set of transition pathways lying in a space of higher dimensionality. Refined approximations of the true MFEP could be obtained by backmapping, or ”uplifting”, the 2D MFEP into a higher-dimensional space provided that the new coordinates offer a better descriptive power of the transition mechanism, that is, if they capture relevant aspects of the transitions overlooked by the coarser description. We developed a method to uplift the 2D eABF-MFEP by embedding it into a 12D CV-space designed to provide a higher-resolution representation of the recovery stroke by 1) including additional degrees of freedom and 2) replacing the coarse-grained Δ*RMSD* Relay helix kink descriptor by 4 separate distance CVs representing local backbone rearrangements upon the formation of the kink. This uplifted string provides the starting point for a new series of CVSM calculations, this time in 12D space.

**Fig S8.**
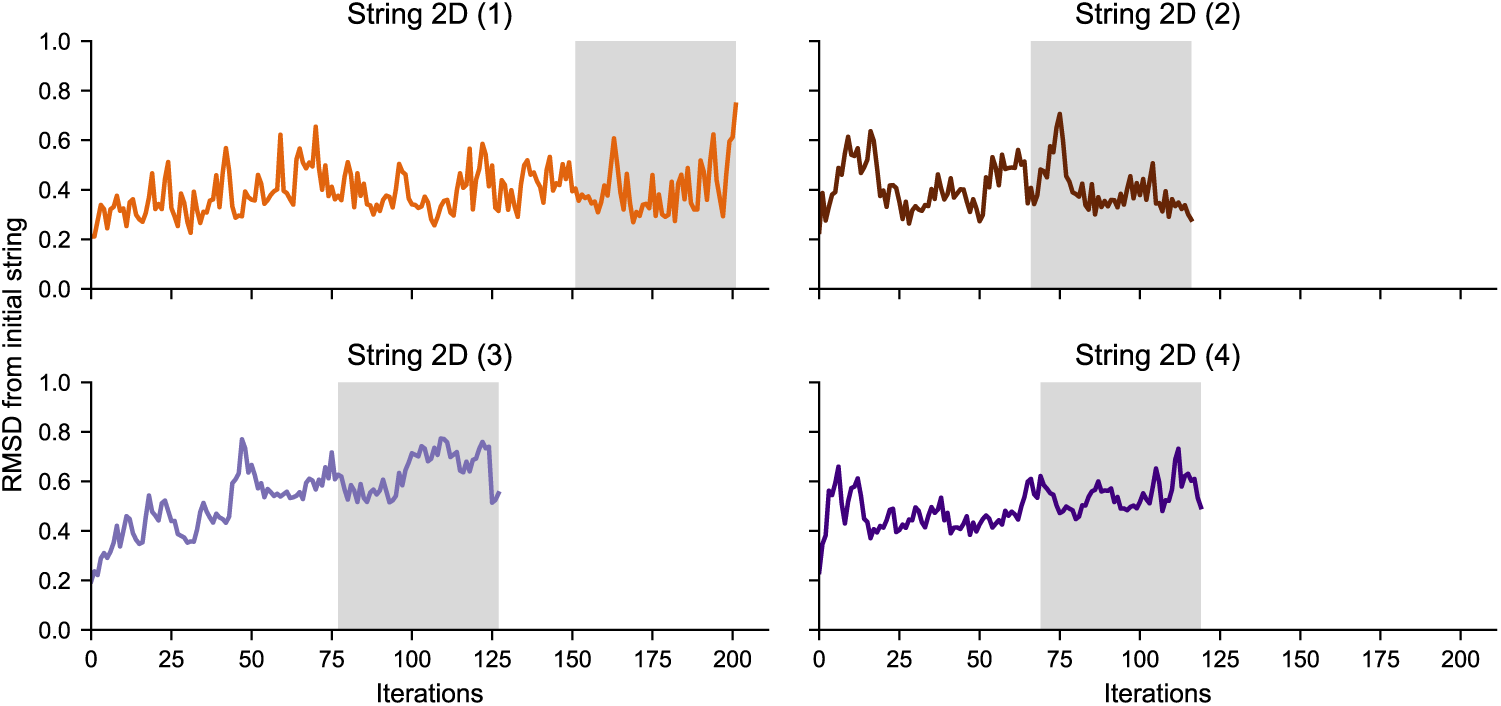
Convergence of CVSM calculations in 2D CV-space. Shown is the RMSD in normalized CVs from the initial path. The greyed area shows the last 50 iterations, over which the average strings pictured in Supplementary Figure S7 are computed.

Such an uplifting requires accounting for the missing information of the additional degrees of freedom. Here, we used the average values of the additional CVs measured from the pre-equilibration Umbrella Sampling simulations along the 2D eABF-MFEP reported above. These provide reasonable initial estimates of the most likely value taken by the additional CVs when the system is restrained in the vicinity of the MFEP. However, the resulting path in 12D CV-space was found to be very rugged, which we reasoned would impede convergence of the string calculations. To alleviate this issue, we regularized the uplifted path by averaging it with a straight path connecting PR to PTS in normalized 12-dimensional CV space. The resulting uplifted regularized path retains the global shape of the uplifted path before regularization, but is much smoother.

##### CVSM calculations from the regularized uplifted string

The 32 Myo6 conformers previously pre-equilibrated in 2D along the eABF-MFEP were further relaxed along the regularized uplifted string in 12D by 1 ns of US. Then, they were used to perform two independent CVSM calculations to reveal the MFEP in 12D CV-space, yielding Strings A1 and A2.

**Table S2.** CVSM simulations in 12D CV space.

| Calculation | Guess path | Ends | $t_{eq}$ (ps) | $n_{swarm}$ | $t_{free}$ (ps) | $n_{iter}$ |
| --- | --- | --- | --- | --- | --- | --- |
| 12D String A1 | uplifted/regularized eABF-MFEP | Fixed | 100 | 20 | 0.5 | 179 |
| 12D String A2 | uplifted/regularized eABF-MFEP | Fixed | 100 | 20 | 0.5 | 119 |
| 12D String B1 | Straight (12D) | Fixed | 100 | 20 | 0.5 | 199 |
| 12D String B2 | Straight (12D) | Fixed | 100 | 20 | 0.5 | 119 |

**Fig S9.**
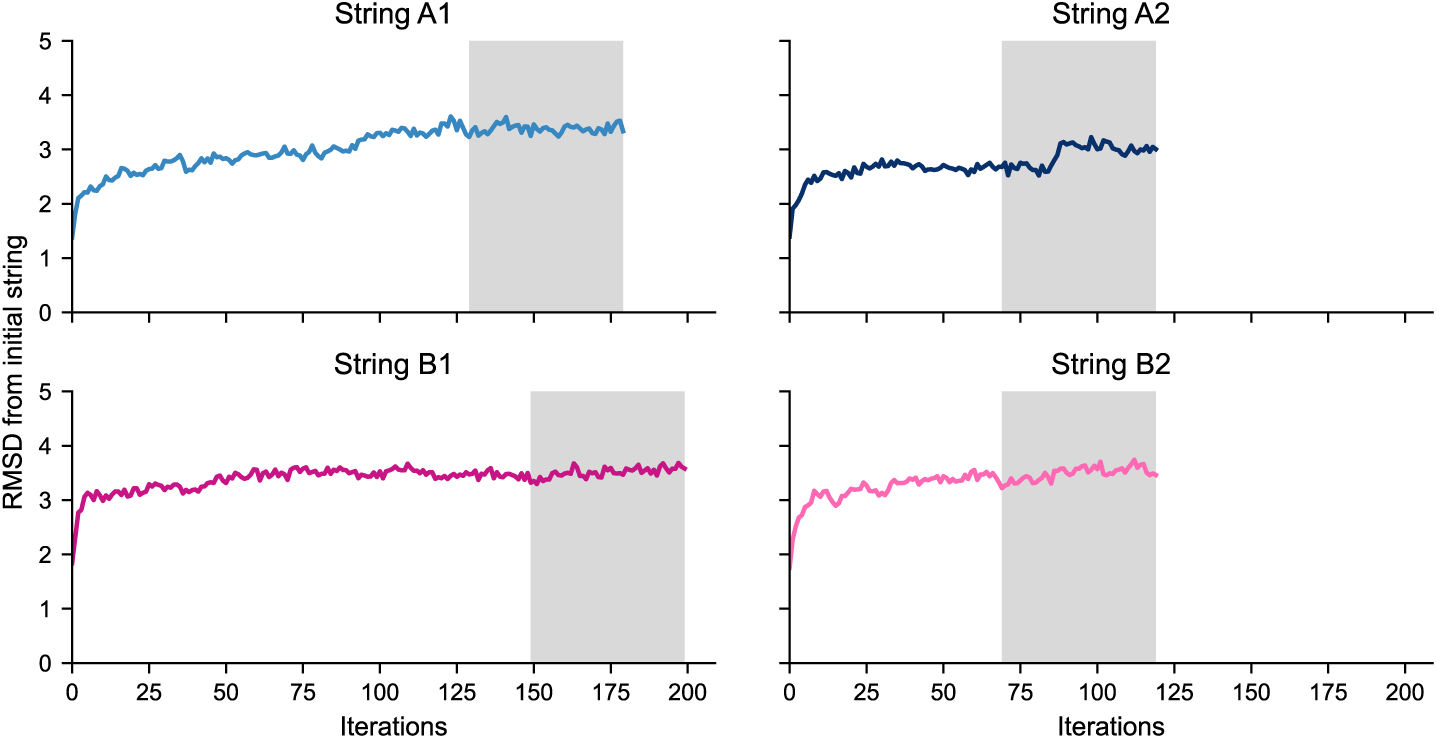
Convergence of CVSM calculations in 12D CV-space. Shown is the RMSD in normalized CVs from the initial path. The greyed area shows the last 50 iterations, over which the average strings are computed.

##### 12D CVSM calculations from the straight guess path

To assess the robustness of the string calculations, we also ran independent string calculations initiated from the linear guess path connecting PR to PTS in normalized 12-dimensional CV-space. 32 equally-spaced conformers along this path were prepared by relaxing the conformers previously pre-equilibrated along the 2D linear path to their corresponding images in the 12D linear string using 1 ns of US. Then, there were used to perform two separate CVSM calculations, yielding Strings B1 and B2. The corresponding transition path, or Path B, is described in Supplementary Text 4.

#### String method calculations in 20D normalized CV-space

The analysis of the transition pathway emerging from the 12D string optimizations initialized with the straight guess path (*i.e.*, Strings B1 and B2) revealed 8 additional potentially relevant degrees of freedom, see Supplementary Text 4. To better reflect their involvement in the mechanism, we uplifted and regularized the converged String B1 to the 20-dimensional CV-space using the procedure described above. The 20D regularized/uplifted string was then relaxed to convergence by 39 iterations of CVSM.

**Fig S10.**
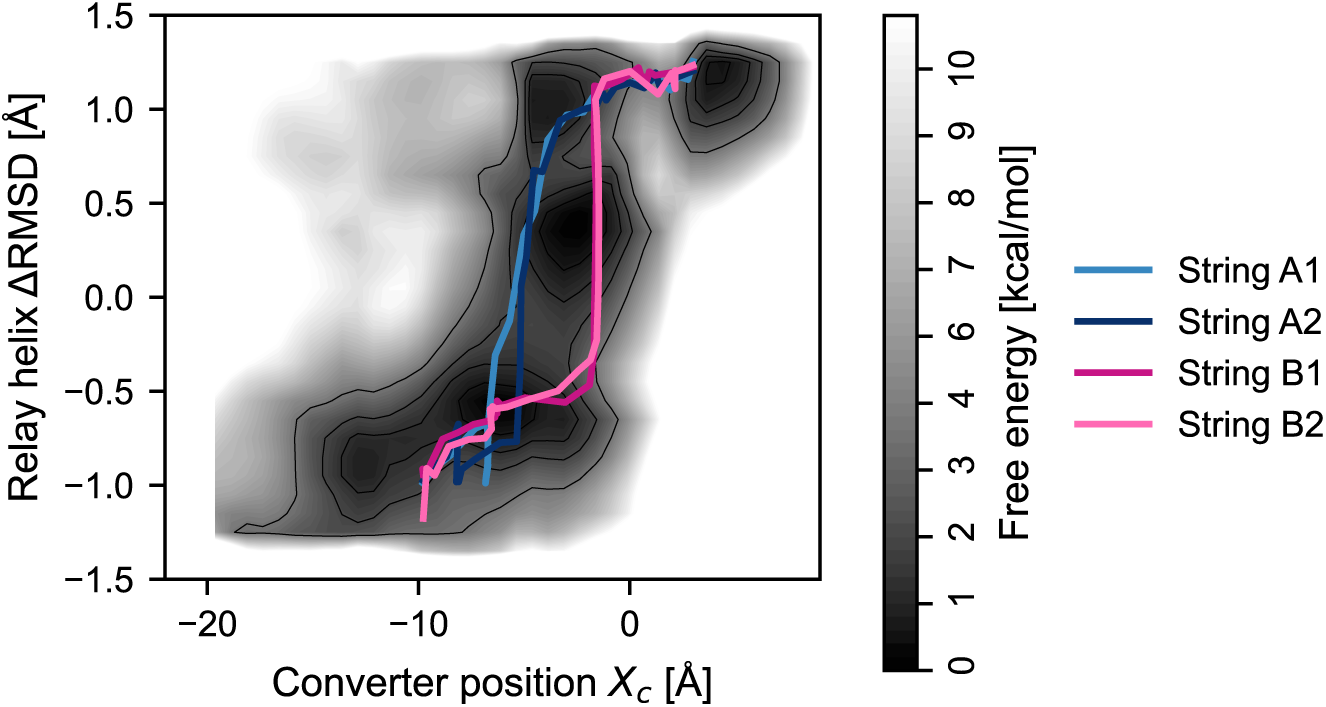
Projections of the 12D converged strings onto 2D space reveal two alternate pathways lying within the transition tube predicted by eABF.

**Fig S11.**
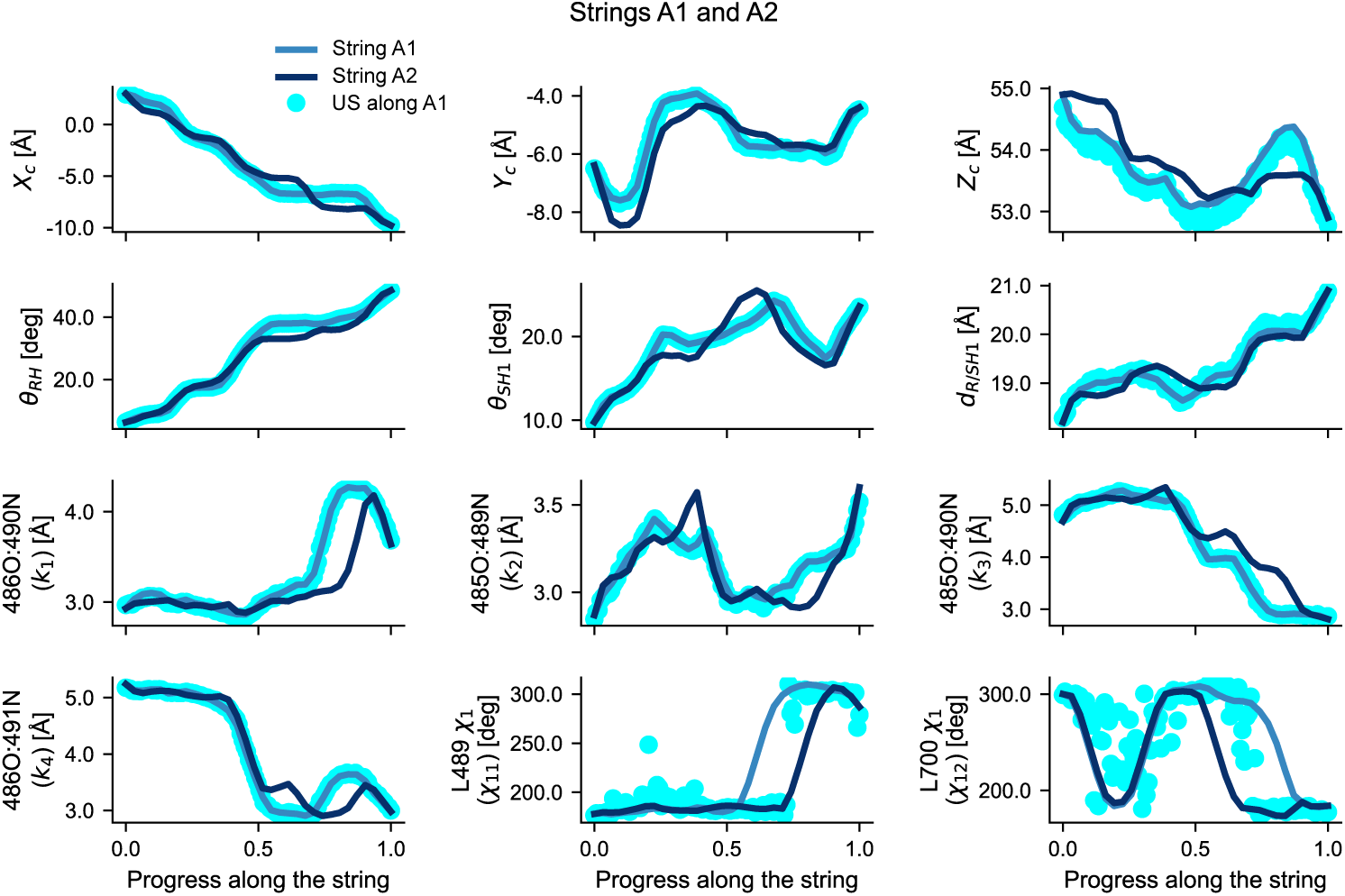
Evolution of biased structural observables along converged, averaged Strings A1 and A2.

**Fig S12.**
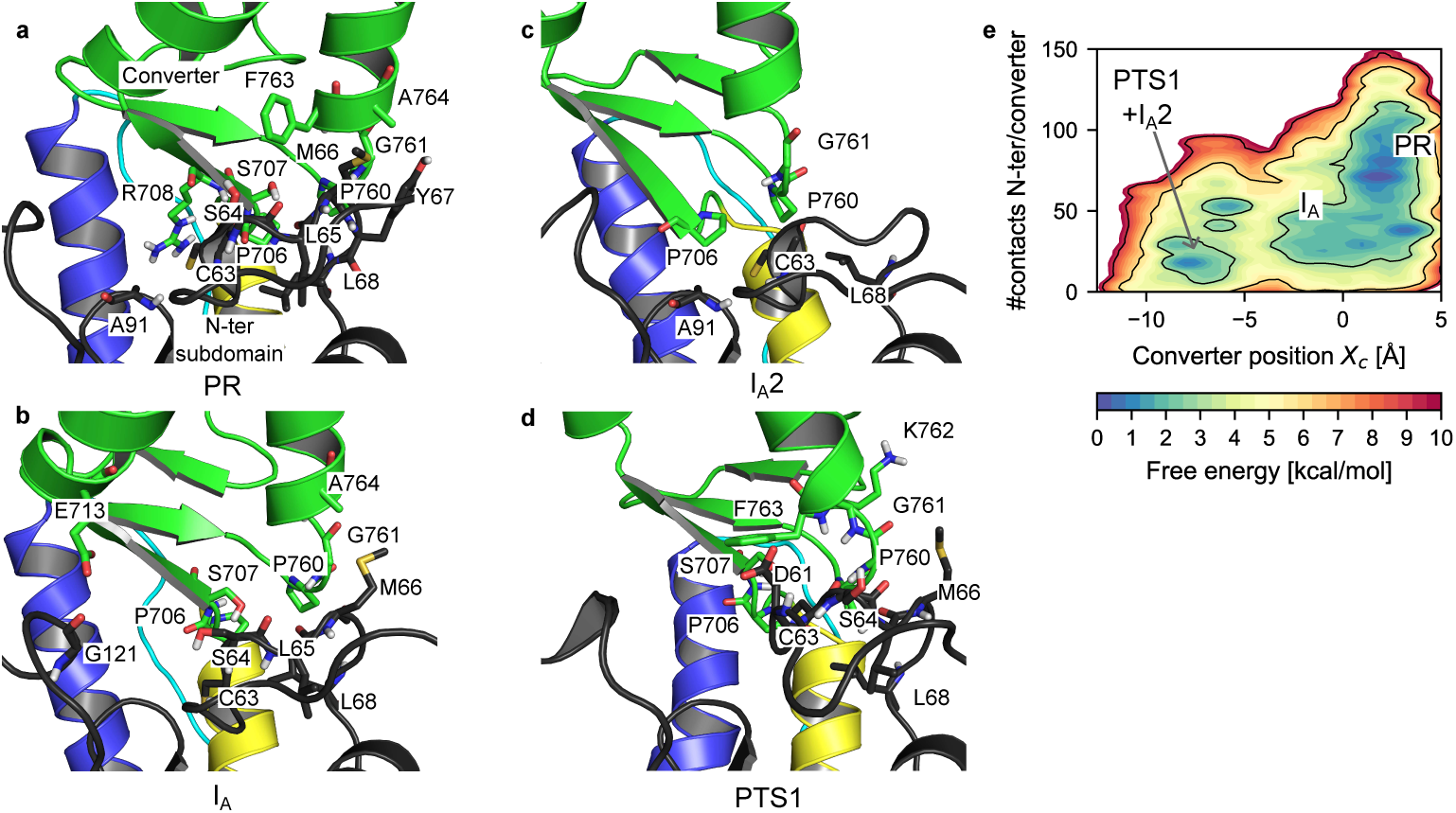
Contacts between the N-terminal subdomain and the converter during the PR→ PTS transition. (a-d) Close-up on the converter/N-ter interface in representative frames of the metastable states sampled in umbrella sampling along String A1. Residues involved in contacts are shown as sticks. For clarity, non-polar hydrogens are not shown. (e) PMF along *X_c_* and the number of contacts estimated with MBAR from umbrella sampling along string A1. A contact is defined as two heavy atoms being closer than 4.5 Å.

**Table S3.**
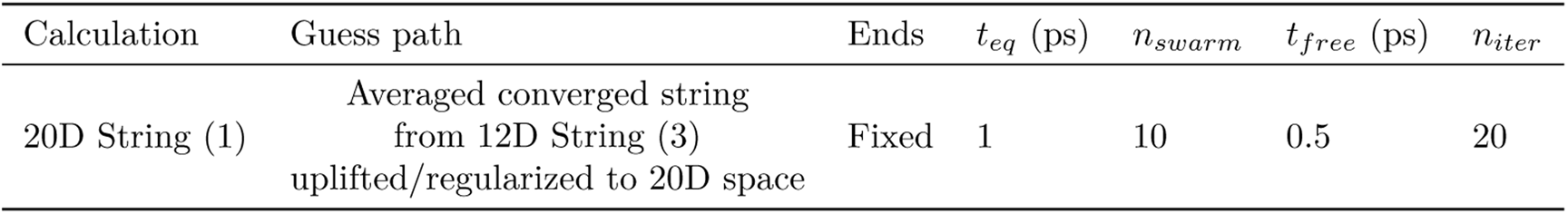
CVSM simulation in 20D CV space.

| Calculation | Guess path | Ends | $t_{eq}$ (ps) | $n_{swarm}$ | $t_{free}$ (ps) | $n_{iter}$ |
| --- | --- | --- | --- | --- | --- | --- |
| 20D String (1) | Averaged converged string<br>from 12D String (3)<br>uplifted/regularized to 20D space | Fixed | 1 | 10 | 0.5 | 20 |

#### Projection of the high-dimensional strings onto 2D (*X*_c_, Δ*RMSD*)-space

We projected the high-dimensional (12 or 20 dimensions) strings onto 2D (*X_c_,* Δ*RMSD*)-space to compare them with the eABF results. Since the Δ*RMSD* CV is not a component of the high-dimensional strings, we evaluated its value as the per-image average over the last 50 string iterations. Since Δ*RMSD* is not explicitly biased nor involved in string reparametrization, the projected strings have a rugged aspect.

**Fig S13.**
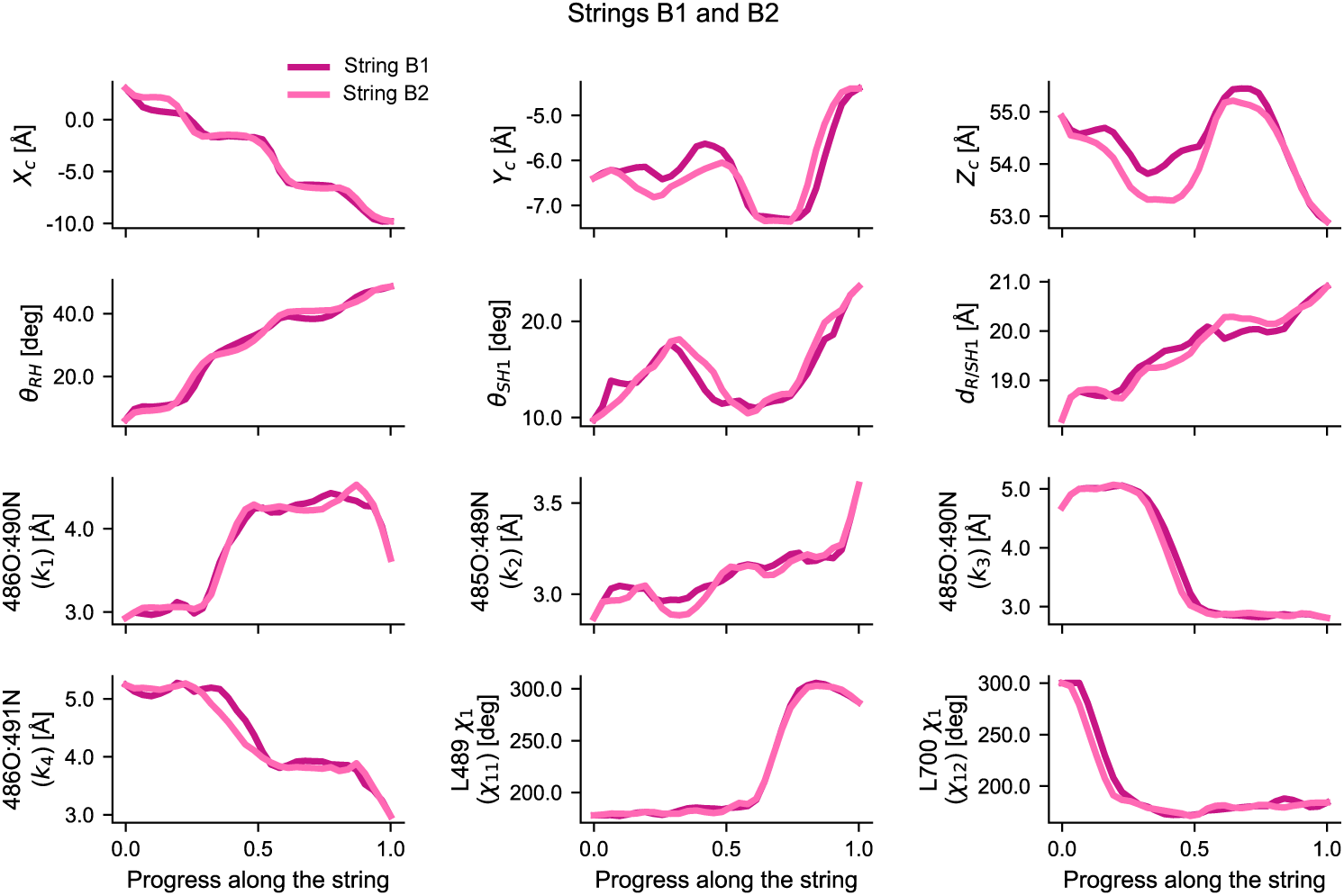
Evolution of biased structural observables along converged, averaged Strings B1 and B2.

**Fig S14.**
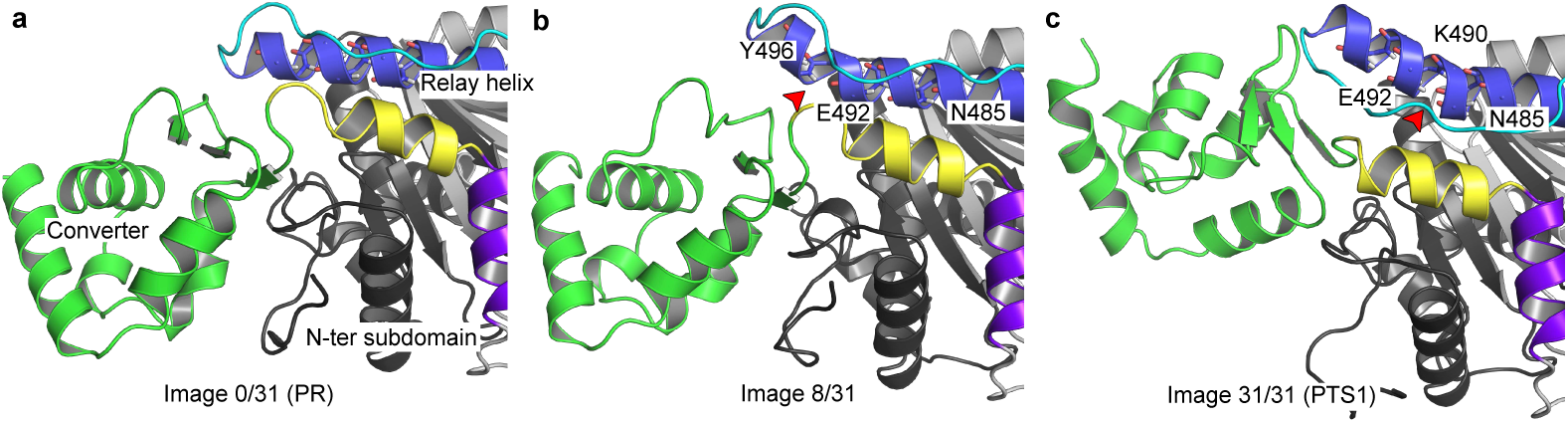
A secondary kink in the Relay helix forms in Path B. a. Initial state (PR-like): the Relay helix is straight and intact, the converter interacts with the N-terminal subdomain. c. *α* =0.25: a secondary kink in the Relay helix (red arrow) accomodates converter movement. The converter is still in contact with the N-terminal subdomain. c. Final state (PTS-like): the converter has moved and broken the contacts, and the canonical kink (red arrow) in the Relay helix has formed.

**Fig S15.**
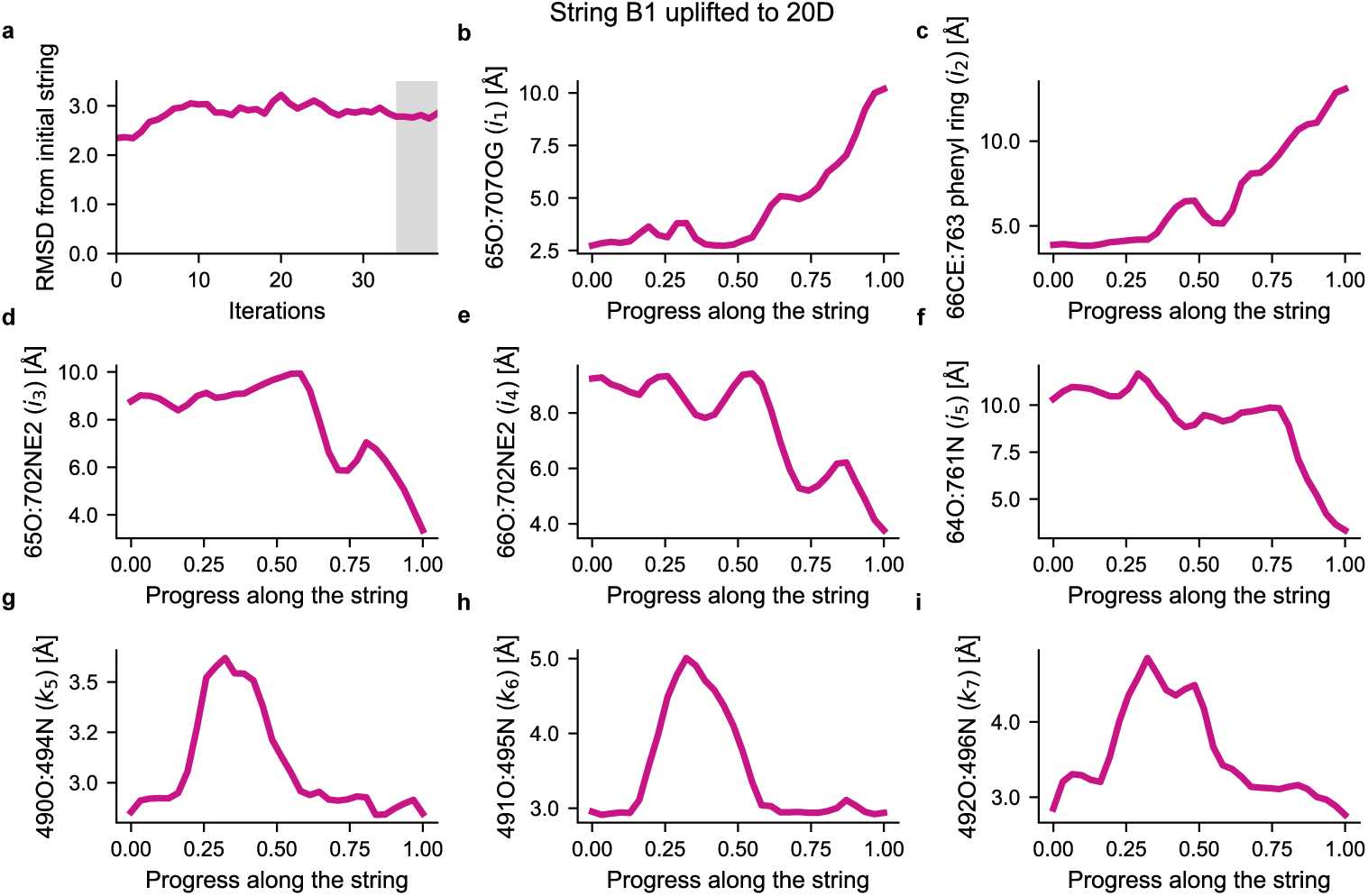
Re-optimization of String B1 in 20-dimensional space. (a) Convergence of the CVSM calculation in 20D CV-space. (b-f) Evolution of converter/N-terminal distances along the converged, averaged 20d string. (g-i) Evolution of the RH backbone distances describing the secondary kink along the converged, averaged 20D string. Panels b and c show that contacts between the converter and the N-terminal subdomain are preserved until about halfway along the transition, while panels g-i show that a secondary kink forms in the RH from about 0.25 to 0.50 progress along the string.

### Supplementary Text 3: free energy profiles along the string from umbrella sampling

#### Umbrella sampling

An ”on-the-path” Umbrella Sampling calculation was performed along the averaged string from the first string optimization initiated from the eABF-regularized guess, or String A1. First, *χ*_11_ and *χ*_12_ were excluded from the set of biased CVs because we reasoned that the sharp nature of rotameric transitions could hinder sampling around the transition state regions. Then, we fitted a collection of *B*-splines on each CV along the string using Scipy. Finally, 128 images equally-spaced along the string in normalized CV space were extracted and used as restraint centers for umbrella sampling. Umbrella sampling simulations were run with GROMACS 2021.5 patched with *colvars*, using the same force constants as for string optimizations, and were initiated from structures sampled in the last iteration of the string optimization. To equilibrate the images with the restraining potentials, the first 1 ns were discarded for each window. Each window was then simulated for 120 ns, resulting in a total of 15.36 µs of umbrella sampling simulation. For analysis, CV data were saved every 10 ps. Free energies along the string were then computed using two different approaches.

#### Free energy profile along the string by umbrella integration

To obtain a PMF along the string, we used the procedure described in [55]. Introducing *α* as a non-dimensional progress variable along the path, such that *α* = 0 at the beginning of the path and *α* = 1 at the end, one can write for the free energy *F* (*α*) along the path:

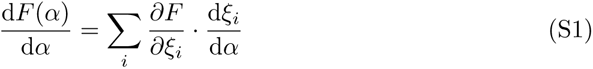

where *ξ_i_* refers to CV *i* along the string, and the sum runs over all supporting CVs. The 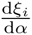 are evaluated by analytical differentiation of the fitted splines. To estimate the free energy gradients with respect to CV components, we used the Umbrella Integration estimator [39] with a number of bins equal to the number of windows, *i.e.* 128.

#### MBAR reweighting

Separately from the UI calculations, free energy profiles along arbitrary CVs (possibly including unrestrained ones) were computed using the Multistate Bennett Acceptance Ratio (MBAR) method [40]. These PMFs represent the effective free energy landscapes when the progress coordinate along the PR → PTS transition is averaged out, and provide insight into the thermodynamic coupling between elementary rearrangements. MBAR calculations were performed using *pymbar* 4.0.1. We used the recently published Kernel Density Estimation (KDE)-based procedure to obtain smooth free energy estimates [41]. In so doing, we found that the choice of the KDE bandwidth parameter could strongly affect the value of the free energy barriers, but only marginally affected the position and relative stability of free energy basins. Therefore, the MBAR-computed free energy maps should be considered semi-quantitative.

### Supplementary Text 4: Mechanism along Path B

From the straight guess path, two independent string optimizations were performed until they converged towards a common sequence of events, which we name Path B (Supplementary Figure S9). Similar to the situation for Path A, Path B from string calculations is virtually identical to the Path *B_ABF_*, as we justify below. Projection of the 2 Path B 12D-strings onto the (*X_c_,* Δ*RMSD*) plane shows that they are consistent with the shape and location of the transition tube (Supplementary Figure S10). Interestingly, Paths A and B do not overlap, and Path B visits I*_B_*. Thus, even if they share the same gross features, the structural mechanism predicted from Path B is a bit different from that of Path A (Supplementary Figure S13). In Path B, the kink in the RH forms with a different timing and mechanism. There, the initial movement of the converter is not accomodated by the disruption of its interactions with the N-terminal subdomain, but by the transient formation of a secondary kink in the RH at the level of residues 490 to 496 (Supplementary Figure S14). This is similar to Path *B_ABF_* from eABF calculations (see notably Fig. 2e) and supports the idea that Paths *B_ABF_* and *B* are in fact representative of the same mechanism. As the hydrogen-bonding distances describing the formation of the secondary kink are not explicitly biased in the 12D string optimizations, it is difficult to know whether the mechanism observed in Path B does correspond to an MFEP also along these degrees of freedom. To clarify this matter, we extended the CV-space to 20 dimensions by adding 3 distances describing the secondary kink and 5 distances describing converter/N-ter interactions, uplifted the first converged Path B string to this space, and relaxed it with the string method (Supplementary Text 2, Supplementary Figure S15a). The resulting pathway still exhibits the same sequence of events, supporting its relevance as a locally optimal MFEP in the PR → PTS transition.

The extended, highly bent configuration of the Relay helix enables the rotation of the converter while some of its contacts with the N-terminal subdomain are preserved (Supplementary Fig. S15b-i, Supplementary Fig. S14). This configuration corresponds to the I*_B_* basin identified in eABF, which may represent a metastable intermediate state along this path. Next, upon going from I*_B_* to PTS1, this secondary kink disappears and the canonical kink in the RH forms concurrently with continued movement of the converter in a fashion similar to Path A.

The sensitivity to the initial guess of the converged path is not overly surprising, as multiple pathways are expected for a system the size of myosin’s even after reducing its dimensionality to 12 (or 20). It is plausible that both families of pathways may actually be explored in the recovery stroke of Myo6. Thus, the Relay helix exhibits significant plasticity throughout its length, which may allow it to bend significantly and even develop transient kinks to accomodate the extensive movement of the converter through a diversity of mechanisms. Nevertheless, Path A is likely to be more representative of the dominant mechanism because it entails less structural perturbation (*i.e.,* only one kink forms), which arguably is a desirable property for an MFEP.

## Notes

### Competing Interest Statement

The authors have declared no competing interest.

### Summary of Updates

Cosmetic improvements to Supplementary Information (no change in scientific content or text)

## References

1. Sweeney HL, Houdusse A. Structural and Functional Insights into the Myosin Motor Mechanism. Annual Review of Biophysics. 2010;39(1):539–557. doi:10.1146/annurev.biophys.050708.133751.

2. Geeves MA, Holmes KC. Structural Mechanism of Muscle Contraction. Annual Review of Biochemistry. 1999;68(1):687–728. doi:10.1146/annurev.biochem.68.1.687.

3. Robert-Paganin J, Pylypenko O, Kikuti C, Sweeney HL, Houdusse A. Force Generation by Myosin Motors: A Structural Perspective. Chemical Reviews. 2020;120(1):5–35. doi:10.1021/acs.chemrev.9b00264.

4. Sweeney HL, Houdusse A. What Can Myosin VI Do in Cells? Current Opinion in Cell Biology. 2007;19(1):57–66. doi:10.1016/j.ceb.2006.12.005.

5. de Jonge JJ, Batters C, O’Loughlin T, Arden SD, Buss F. The MYO6 Interactome: Selective Motor-Cargo Complexes for Diverse Cellular Processes. FEBS Letters. 2019;593(13):1494–1507. doi:10.1002/1873-3468.13486.

6. Ménétrey J, Bahloul A, Wells AL, Yengo CM, Morris CA, Sweeney HL, et al. The Structure of the Myosin VI Motor Reveals the Mechanism of Directionality Reversal. Nature. 2005;435(7043):779–785. doi:10.1038/nature03592.

7. Ménétrey J, Llinas P, Mukherjea M, Sweeney HL, Houdusse A. The Structural Basis for the Large Powerstroke of Myosin VI. Cell. 2007;131(2):300–308. doi:10.1016/j.cell.2007.08.027.

8. Ménétrey J, Llinas P, Cicolari J, Squires G, Liu X, Li A, et al. The Post-Rigor Structure of Myosin VI and Implications for the Recovery Stroke. The EMBO Journal. 2008;27(1):244–252. doi:10.1038/sj.emboj.7601937.

9. Fisher AJ, Smith CA, Thoden J, Smith R, Sutoh K, Holden HM, et al. X-Ray Structures of the Myosin Motor Domain of Dictyostelium Discoideum Complexed with MgADP.BeFx and MgADP.AlF4-. Biochemistry. 1995;34(28):8960–8972. doi:10.1021/bi00028a004.

10. Kiani FA, Fischer S. Catalytic Strategy Used by the Myosin Motor to Hydrolyze ATP. Proceedings of the National Academy of Sciences. 2014;111(29):E2947–E2956. doi:10.1073/pnas.1401862111.

11. Kiani FA, Fischer S. Stabilization of the ADP/Metaphosphate Intermediate during ATP Hydrolysis in Pre-power Stroke Myosin: QUANTITATIVE ANATOMY OF AN ENZYME. Journal of Biological Chemistry. 2013;288(49):35569–35580. doi:10.1074/jbc.M113.500298.

12. Lu X, Ovchinnikov V, Demapan D, Roston D, Cui Q. Regulation and Plasticity of Catalysis in Enzymes: Insights from Analysis of Mechanochemical Coupling in Myosin. Biochemistry. 2017;56(10):1482–1497. doi:10.1021/acs.biochem.7b00016.

13. Geeves MA. Review: The ATPase Mechanism of Myosin and Actomyosin: The ATPase Mechanism of Myosin and Actomyosin. Biopolymers. 2016;105(8):483–491. doi:10.1002/bip.22853.

14. Fischer S, Windshügel B, Horak D, Holmes KC, Smith JC. Structural Mechanism of the Recovery Stroke in the Myosin Molecular Motor. Proceedings of the National Academy of Sciences of the United States of America. 2005;102(19):6873–6878. doi:10.1073/pnas.0408784102.

15. Koppole S, Smith JC, Fischer S. Simulations of the Myosin II Motor Reveal a Nucleotide-state Sensing Element That Controls the Recovery Stroke. Journal of Molecular Biology. 2006;361(3):604–616. doi:10.1016/j.jmb.2006.06.022.

16. Mesentean S, Koppole S, Smith JC, Fischer S. The Principal Motions Involved in the Coupling Mechanism of the Recovery Stroke of the Myosin Motor. Journal of Molecular Biology. 2007;367(2):591–602. doi:10.1016/j.jmb.2006.12.058.

17. Koppole S, Smith JC, Fischer S. The Structural Coupling between ATPase Activation and Recovery Stroke in the Myosin II Motor. Structure. 2007;15(7):825–837. doi:10.1016/j.str.2007.06.008.

18. Woo HJ. Exploration of the Conformational Space of Myosin Recovery Stroke via Molecular Dynamics. Biophysical Chemistry. 2007;125(1):127–137. doi:10.1016/j.bpc.2006.07.001.

19. Harris MJ, Woo HJ. Energetics of Subdomain Movements and Fluorescence Probe Solvation Environment Change in ATP-bound Myosin. European Biophysics Journal. 2008;38(1):1–12. doi:10.1007/s00249-008-0347-3.

20. Yu H, Ma L, Yang Y, Cui Q. Mechanochemical Coupling in the Myosin Motor Domain. I. Insights from Equilibrium Active-Site Simulations. PLoS Computational Biology. 2007;3(2):e21. doi:10.1371/journal.pcbi.0030021.

21. Yu H, Ma L, Yang Y, Cui Q. Mechanochemical Coupling in the Myosin Motor Domain. II. Analysis of Critical Residues. PLoS Computational Biology. 2007;3(2):e23. doi:10.1371/journal.pcbi.0030023.

22. Elber R, West A. Atomically Detailed Simulation of the Recovery Stroke in Myosin by Milestoning. Proceedings of the National Academy of Sciences. 2010;107(11):5001–5005. doi:10.1073/pnas.0909636107.

23. Baumketner A, Nesmelov Y. Early Stages of the Recovery Stroke in Myosin II Studied by Molecular Dynamics Simulations. Protein Science. 2011;20(12):2013–2022. doi:10.1002/pro.737.

24. Baumketner A. Interactions between Relay Helix and Src Homology 1 (SH1) Domain Helix Drive the Converter Domain Rotation during the Recovery Stroke of Myosin II. Proteins: Structure, Function, and Bioinformatics. 2012;80(6):1569–1581. doi:10.1002/prot.24051.

25. Baumketner A. The Mechanism of the Converter Domain Rotation in the Recovery Stroke of Myosin Motor Protein. Proteins: Structure, Function, and Bioinformatics. 2012;80(12):2701–2710. doi:10.1002/prot.24155.

26. Blanc F, Isabet T, Benisty H, Sweeney HL, Cecchini M, Houdusse A. An Intermediate Along the Recovery Stroke of Myosin VI Revealed by X-ray Crystallography and Molecular Dynamics. Proceedings of the National Academy of Sciences. 2018; p. 201711512. doi:10.1073/pnas.1711512115.

27. Torrie GM, Valleau JP. Nonphysical Sampling Distributions in Monte Carlo Free-Energy Estimation: Umbrella Sampling. Journal of Computational Physics. 1977;23(2):187–199. doi:10.1016/0021-9991(77)90121-8.

28. Laio A, Parrinello M. Escaping Free-Energy Minima. Proceedings of the National Academy of Sciences. 2002;99(20):12562–12566. doi:10.1073/pnas.202427399.

29. Comer J, Gumbart JC, Hénin J, Lelièvre T, Pohorille A, Chipot C. The Adaptive Biasing Force Method: Everything You Always Wanted To Know but Were Afraid To Ask. The Journal of Physical Chemistry B. 2015;119(3):1129–1151. doi:10.1021/jp506633n.

30. Chipot C, Pohorille A, editors. Free Energy Calculations: Theory and Applications in Chemistry and Biology. No. 86 in Springer Series in Chemical Physics. Berlin ; New York: Springer; 2007.

31. Maragliano L, Fischer A, Vanden-Eijnden E, Ciccotti G. String Method in Collective Variables: Minimum Free Energy Paths and Isocommittor Surfaces. The Journal of Chemical Physics. 2006;125(2):024106. doi:10.1063/1.2212942.

32. Pan AC, Sezer D, Roux B. Finding Transition Pathways Using the String Method with Swarms of Trajectories. The Journal of Physical Chemistry B. 2008;112(11):3432–3440. doi:10.1021/jp0777059.

33. Ma W, Schulten K. Mechanism of Substrate Translocation by a Ring-Shaped ATPase Motor at Millisecond Resolution. Journal of the American Chemical Society. 2015;137(8):3031–3040. doi:10.1021/ja512605w.

34. Lev B, Murail S, Poitevin F, Cromer BA, Baaden M, Delarue M, et al. String Method Solution of the Gating Pathways for a Pentameric Ligand-Gated Ion Channel. Proceedings of the National Academy of Sciences. 2017;114(21):E4158–E4167. doi:10.1073/pnas.1617567114.

35. Singharoy A, Chipot C, Moradi M, Schulten K. Chemomechanical Coupling in Hexameric Protein–Protein Interfaces Harnesses Energy within V-Type ATPases. Journal of the American Chemical Society. 2017;139(1):293–310. doi:10.1021/jacs.6b10744.

36. Ovchinnikov V, Karplus M, Vanden-Eijnden E. Free Energy of Conformational Transition Paths in Biomolecules: The String Method and Its Application to Myosin VI. The Journal of Chemical Physics. 2011;134(8):085103. doi:10.1063/1.3544209.

37. Ovchinnikov V, Cecchini M, Vanden-Eijnden E, Karplus M. A Conformational Transition in the Myosin VI Converter Contributes to the Variable Step Size. Biophysical Journal. 2011;101(10):2436–2444. doi:10.1016/j.bpj.2011.09.044.

38. Lesage A, Lelièvre T, Stoltz G, Hénin J. Smoothed Biasing Forces Yield Unbiased Free Energies with the Extended-System Adaptive Biasing Force Method. The Journal of Physical Chemistry B. 2017;121(15):3676–3685. doi:10.1021/acs.jpcb.6b10055.

39. Kästner J, Thiel W. Bridging the Gap Between Thermodynamic Integration and Umbrella Sampling Provides a Novel Analysis Method: “Umbrella Integration”. The Journal of Chemical Physics. 2005;123(14):144104. doi:10.1063/1.2052648.

40. Shirts MR, Chodera JD. Statistically Optimal Analysis of Samples from Multiple Equilibrium States. The Journal of Chemical Physics. 2008;129(12):124105. doi:10.1063/1.2978177.

41. Shirts MR, Ferguson AL. Statistically Optimal Continuous Free Energy Surfaces from Biased Simulations and Multistate Reweighting. Journal of Chemical Theory and Computation. 2020;16(7):4107–4125. doi:10.1021/acs.jctc.0c00077.

42. Hill TL. Free Energy Transduction and Biochemical Cycle Kinetics. Mineola, N.Y: Dover Publications; 2005.

43. Sirigu S, Hartman JJ, Planelles-Herrero VJ, Ropars V, Clancy S, Wang X, et al. Highly Selective Inhibition of Myosin Motors Provides the Basis of Potential Therapeutic Application. Proceedings of the National Academy of Sciences. 2016; p. 201609342. doi:10.1073/pnas.1609342113.

44. Akter F, Ochala J, Fornili A. Binding Pocket Dynamics along the Recovery Stroke of Human *β*-Cardiac Myosin. PLOS Computational Biology. 2023;19(5):e1011099. doi:10.1371/journal.pcbi.1011099.

45. Yang Y, Yu H, Cui Q. Extensive Conformational Transitions Are Required to Turn On ATP Hydrolysis in Myosin. Journal of Molecular Biology. 2008;381(5):1407–1420. doi:10.1016/j.jmb.2008.06.071.

46. Baldo AP, Tardiff JC, Schwartz SD. Mechanochemical Function of Myosin II: Investigation into the Recovery Stroke and ATP Hydrolysis. The Journal of Physical Chemistry B. 2020;124(45):10014–10023. doi:10.1021/acs.jpcb.0c05762.

47. Chakraborti A, Baldo AP, Tardiff JC, Schwartz SD. Investigation of the Recovery Stroke and ATP Hydrolysis and Changes Caused Due to the Cardiomyopathic Point Mutations in Human Cardiac *β* Myosin. The Journal of Physical Chemistry B. 2021; p. acs.jpcb.1c03144. doi:10.1021/acs.jpcb.1c03144.

48. Porter JR, Meller A, Zimmerman MI, Greenberg MJ, Bowman GR. Conformational Distributions of Isolated Myosin Motor Domains Encode Their Mechanochemical Properties. eLife. 2020;9:e55132. doi:10.7554/eLife.55132.

49. Meller A, Lotthammer JM, Smith LG, Novak B, Lee LA, Kuhn CC, et al. Drug Specificity and Affinity Are Encoded in the Probability of Cryptic Pocket Opening in Myosin Motor Domains. eLife. 2023;12:e83602. doi:10.7554/eLife.83602.

50. Chakraborti A, Tardiff JC, Schwartz SD. Insights into the Mechanism of the Cardiac Drug Omecamtiv Mecarbil-A Computational Study. The Journal of Physical Chemistry B. 2022;doi:10.1021/acs.jpcb.2c06679.

51. Phillips JC, Braun R, Wang W, Gumbart J, Tajkhorshid E, Villa E, et al. Scalable Molecular Dynamics with NAMD. Journal of Computational Chemistry. 2005;26(16):1781–1802. doi:10.1002/jcc.20289.

52. Abraham MJ, Murtola T, Schulz R, Páll S, Smith JC, Hess B, et al. GROMACS: High Performance Molecular Simulations through Multi-Level Parallelism from Laptops to Supercomputers. SoftwareX. 2015;1–2:19–25. doi:10.1016/j.softx.2015.06.001.

53. Fiorin G, Klein ML, Hénin J. Using Collective Variables to Drive Molecular Dynamics Simulations. Molecular Physics. 2013;111(22-23):3345–3362. doi:10.1080/00268976.2013.813594.

54. Hénin J, Chipot C. Overcoming Free Energy Barriers Using Unconstrained Molecular Dynamics Simulations. The Journal of Chemical Physics. 2004;121(7):2904. doi:10.1063/1.1773132.

55. Blanc FE, Cecchini M. An Asymmetric Mechanism in a Symmetric Molecular Machine. The Journal of Physical Chemistry Letters. 2021;12(13):3260–3265. doi:10.1021/acs.jpclett.1c00404.

56. Johnson ME, Hummer G. Characterization of a Dynamic String Method for the Construction of Transition Pathways in Molecular Reactions. The Journal of Physical Chemistry B. 2012;116(29):8573–8583. doi:10.1021/jp212611k.

57. Maragliano L, Roux B, Vanden-Eijnden E. Comparison between Mean Forces and Swarms-of-Trajectories String Methods. Journal of Chemical Theory and Computation. 2014;10(2):524–533. doi:10.1021/ct400606c.

58. Pedregosa F, Varoquaux G, Gramfort A, Michel V, Thirion B, Grisel O, et al. Scikit-Learn: Machine Learning in Python. MACHINE LEARNING IN PYTHON. 2011; p. 6.

59. Wereszczynski J, McCammon JA. Nucleotide-Dependent Mechanism of Get3 as Elucidated from Free Energy Calculations. Proceedings of the National Academy of Sciences. 2012;109(20):7759–7764.

60. E W, Ren W, Vanden-Eijnden E. String Method for the Study of Rare Events. Physical Review B. 2002;66(5). doi:10.1103/PhysRevB.66.052301.

61. Jones E, Oliphant T, Peterson P. {SciPy}: Open Source Scientific Tools for {Python}. 2001–///;.

62. Virtanen P, Gommers R, Oliphant TE, Haberland M, Reddy T, Cournapeau D, et al. SciPy 1.0: Fundamental Algorithms for Scientific Computing in Python. Nature Methods. 2020;17(3):261–272. doi:10.1038/s41592-019-0686-2.

